# Learning the shape of protein micro-environments with a holographic convolutional neural network

**DOI:** 10.1101/2022.10.31.514614

**Authors:** Michael N. Pun, Andrew Ivanov, Quinn Bellamy, Zachary Montague, Colin LaMont, Philip Bradley, Jakub Otwinowski, Armita Nourmohammad

## Abstract

Proteins play a central role in biology from immune recognition to brain activity. While major advances in machine learning have improved our ability to predict protein structure from sequence, determining protein function from structure remains a major challenge. Here, we introduce Holographic Convolutional Neural Network (H-CNN) for proteins, which is a physically motivated machine learning approach to model amino acid preferences in protein structures. H-CNN reflects physical interactions in a protein structure and recapitulates the functional information stored in evolutionary data. H-CNN accurately predicts the impact of mutations on protein function, including stability and binding of protein complexes. Our interpretable computational model for protein structure-function maps could guide design of novel proteins with desired function.

---

Proteins are the machinery of life. They facilitate the key processes that drive living organisms and only rely on twenty types of amino acids to do so. The physical arrangement of a protein’s amino acids dictates how it folds and interacts with its environment. While this compositional nature gives rise to the diversity of existing proteins, it also makes determining protein function from structure a complex problem.

With the growing amount of data and computational advances, machine learning has come to the forefront of protein science, especially in predicting structure from sequence [1–5]. However, the problem of how protein function is determined from its sequence or structure still remains a major challenge.

Techniques from natural language processing are used to determine functional motifs in protein sequences by allowing residues far away in sequence to form information units about function [6–12]. However, since protein function is closely related to protein structure, models trained to map protein sequences to function must account for the complex sequence-structure map.

Despite AlphaFold’s remarkable success at predicting protein folding, it still struggles to determine the effect of mutations on the stability and function of a protein [13]. Nonetheless, it is suggested that AlphaFold has learned an effective physical potential to fold proteins, and therefore, it could be used to characterize the effect of mutations or general protein function [14]. Given the availability of high-resolution tertiary structures, obtained either experimentally or computationally, the information on atomic coordinates in a 3D structure of a protein can be used to learn a direct map to function. Indeed, models aware of the geometry of protein 3D structure that attempt to solve the inverse folding problem, i.e., designing a sequence that folds into a desired structure, can be used to reliably infer the functional effect of mutations in a protein sequence [15], or even engineer diverse sequences that have a desired function [16].

Recent work in the field of molecular dynamics (MD) has shown the power of geometry-aware machine learning at inferring precise inter-atomic force fields [17–24]. Compared to the geometry-aware protein structure models [3, 4, 15, 16], the MD models use more complex geometric features, resulting in more expressive, yet physically interpretable models of molecular interactions.

Here, we introduce holographic convolutional neural network (H-CNN) to learn physically grounded structure-to-function maps for proteins. H-CNN learns local representations of protein structures by encoding the atoms within protein micro-environments in a spherical Fourier space as holograms, and processes these holographic projections via a 3D rotationally equivariant convolutional neural network (Fig. 1) [25–27]. The resulting model respects rotational symmetry of protein structures and characterizes effective inter-amino-acid potentials in protein micro-environments.

**FIG. 1.**
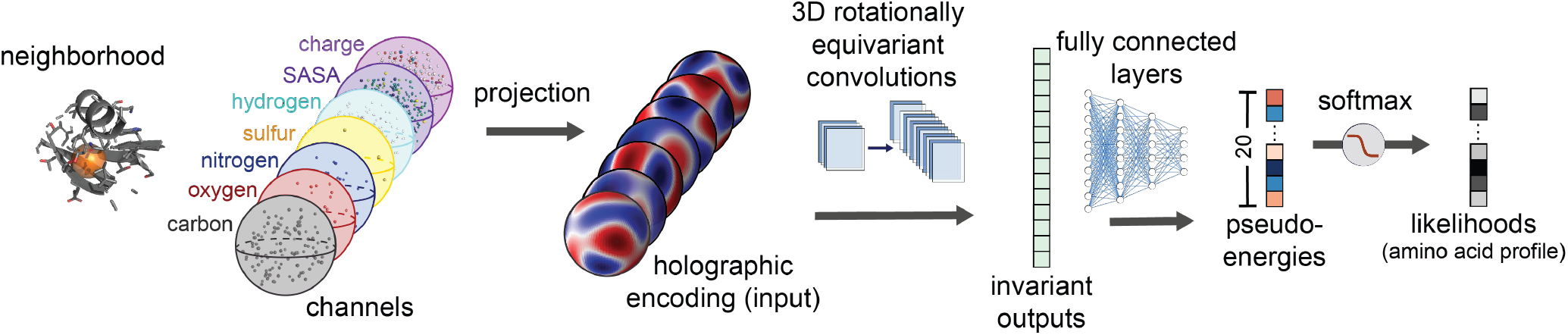
Schematic of Holographic Convolutional Neural network (H-CNN) for protein micro-environments. A neighborhood within a radius of 10Å around a focal amino acid (masked in orange) in a protein structure is separated into its constituent atomic and chemical channels. The information in these channels is encoded in a rotationally equivariant form, using 3D Zernike polynomials, which defines holograms in spherical Fourier space. These holographic encodings are processed by a rotationally equivariant convolutional neural network (Clebsch-Gordan net [26]). The invariant features of the network layers are then collected and processed through fully connected feed-forward layers to determine the preferences, i.e., statistical weights (pseudo-energies) and probabilities, for different amino acids residing at the center of the input neighborhood. The set of predicted probability vectors across all 20 amino acids defines an amino acid profile. The network is trained by learning the categorical classification task with a softmax cross-entropy loss on a one-hot label of the neighborhood determined by the true central residue in the protein structure. A more detailed network architecture is presented in Fig. S2.

We train H-CNN on protein structures available in the Protein Data Bank (PDB) [28], and perform the supervised task of predicting the identity of an amino acid from its surrounding atomic neighborhood with a high accuracy and computational efficiency. The amino acids that H-CNN infers to be interchangeable have similar physicochemical properties, and the pattern is consistent with substitution patterns in evolutionary data. The H-CNN model encodes a more complete set of geometric features of protein structures compared to the other geometry-aware models of proteins [12, 15, 16]. Therefore, it can predict protein function, including binding and stability effects of mutations, only based on local atomic compositions in a protein structure. Our results showcase that principled geometry-aware machine learning can lead to powerful and robust models that provide insight into the biophysics of protein interactions and function, with a potential for protein design.

## I. RESULTS

### Model

We define the micro-environment surrounding an amino acid as the identity and the 3D coordinates of atoms within a radius of 10 Å of the focal amino acid’s α-carbon; this neighborhood excludes atoms from the focal amino acid.

A common approach to encode such atomic neighborhoods for computational analysis is to voxelize the coordinates, which is a form of binning in 3D [30, 31]. However, this approach distorts the information, since the voxel boundaries are arbitrary—too large voxels average over many atoms, and too small voxels lead to very sparse data.

The other obstacle for modeling such data is more fundamental and related to the rotational symmetries in encoding a protein structure neighborhood. A given neighborhood can occur in different orientations within or across proteins, and a machine learning algorithm should account for such rotational symmetry. One approach known as data augmentation, mainly used in image processing, trains an algorithm on many examples of an image in different orientations and locations. Data augmentation is computationally costly in 3D, and it is likely to result in a model of amino acid interactions that depends on the neighborhood’s orientation, which is a non-physical outcome. Another approach is to orient the amino acid neighborhoods based on a prior choice (e.g., along the backbone of the protein) [30, 31]. However, this choice is somewhat arbitrary, and the specified orientation of the protein backbone could inform the model about the identity of the focal amino acid.

To overcome these obstacles, we introduce holographic convolutional neural networks (H-CNNs) for protein micro-environments. First, we encode the point clouds of different atoms in an amino acid neighborhood using 3D Zernike polynomials as spherical basis functions (Fig. 1 and Methods). 3D Zernike polynomials can be used to expand any function in three dimensions along angular and radial bases and can uniquely represent the properties of the encoded object in a spherical Fourier space, given enough terms in the Fourier series. Conveniently, the angular component of the Zernike polynomials are spherical harmonics, which form an equivariant basis under rotation in 3D. Rotational equivariance is the property that if the input (i.e., atomic coordinates of an amino acid’s neighborhood) is rotated, then the output is transformed according to a linear operation determined by the rotation (Fig. S1). As a result, these Zernike projections enable us to encode the atomic point clouds from a protein structure without having to align the neighborhoods. Zernike projections in spherical Fourier space can be understood as a superposition of spherical holograms of an input point cloud, and thus, we term this operation as *holographic encoding* of protein micro-environments; see Fig. 1 and Fig. S2 and Methods for details.

The holograms encoding protein neighborhoods are input to a type of convolutional neural network (CNN). This network is trained on the supervised task of predicting the identity of a focal amino acid from the surrounding atoms in the protein’s tertiary structure. Conventional CNNs average over spatial translations and can learn features in the data that may be in different locations (i.e., they respect translational symmetry). For the analysis of protein neighborhoods we need to infer models that are insensitive to the orientation of the data (i.e., that respect 3D rotational symmetry of the point clouds in a protein neighborhood). Recent work in machine learning has expanded CNNs to respect physical symmetries beyond translations [25–27]. For 3D rotations, generalized convolutions use spherical harmonics, which arise from the irreducible representations of the 3D rotation group SO(3) [32]. For our analysis, we use Clebsch-Gordon neural networks [26], in which the linear and the nonlinear operations of the network layers have the property of rotational equivariance (Methods); see Fig. S2 for detailed information on network architecture and Fig. S3 and Table S1 for details on hyper-parameter tuning and training of the network.

Taken together, the H-CNN shown in Figs. 1, S2, takes as input holograms that encode the spatial composition of different atoms (Carbon, Nitrogen, Oxygen, Sulfur, Hydrogen) and physical properties such as charge and solvent accessible surface area (SASA). The input is processed by a 3D rotationally equivariant CNN to learn statistical representations for protein neighborhoods. We train this H-CNN as a classifier on protein neighborhoods, collected from tertiary structures from the PDB, and use the trained network to quantify the preferences for different amino acids in a given structural neighborhood; see Methods for details on data pre-processing.

### H-CNN reveals physico-chemical properties of amino acids, consistent with evolutionary variation

H-CNN predicts the identity of an amino acid from its surrounding micro-environment with 68% accuracy (Fig. 2). The accuracy of H-CNN is comparable to state-of-the-art approaches with conventional CNNs that voxelize and orient the data along the backbone of a central amino acid, while using a smaller atomic micro-environement for performing this classification task [30, 31]; see Table S2 for a detailed comparison of models. Notably, restricting the training of H-CNN to the subspace of models that are rotationally equivariant leads to a substantial speedup in the training of H-CNN compared to the conventional techniques [30, 31]. Moreover, H-CNN is more accurate than other symmetry-aware approaches for molecular modeling, while using an order of magnitude fewer parameters [33, 34]; see Methods and Table S2 for a detailed comparison of models.

**FIG. 2.**
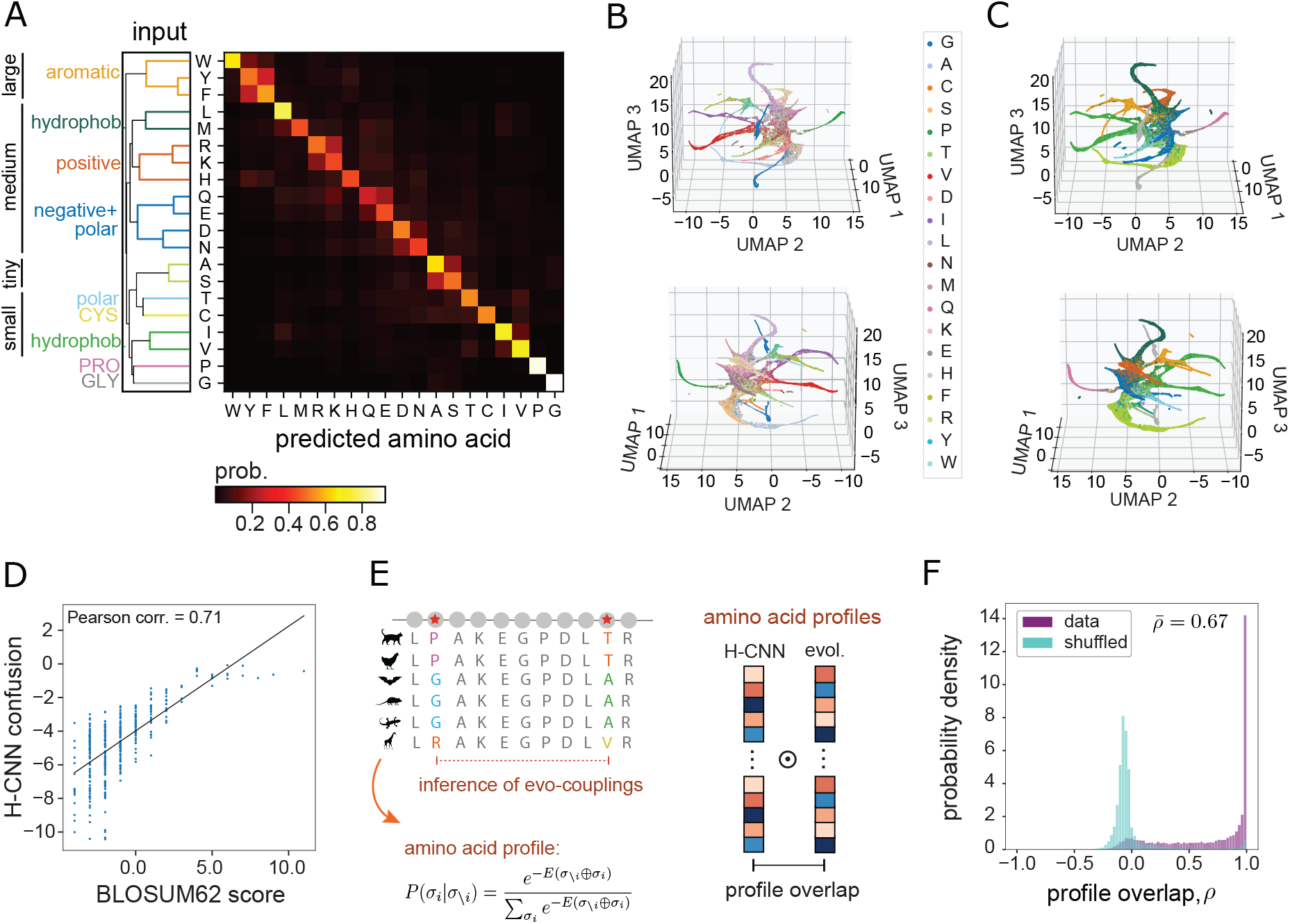
H-CNN predicts amino acid preferences in protein micro-environments. **(A)** The confusion matrix for amino acid predictions with H-CNN shows the mean H-CNN predicted probabilities of each of the twenty amino acids (output) conditioned on a specific central amino acid (input). Overall prediction accuracy is 68%. The hierarchical clustering for these predictions reflects known similarities in size and physico-chemical properties of amino acids. **(B,C)** Low dimensional projections of the prediction outputs (3D UMAPs) are shown. UMAPs are annotated by **(B)** the amino acid types, and **(C)** the physico-chemical clusters in (A), with panels showing a different view of the UMAP in each case. Neighborhoods are closely clustered by amino acid types (B), and are spatially arranged based on the physico-chemical properties of the side-chains (C); colors in (C) are consistent with (A). **(D)** Amino acid confusion in (A) correlates with the substitutability of amino acids in natural proteins as determined by the BLOSUM62 matrix; 71% Pearson correlation. **(E)** Schematic shows how evolutionary covariation of amino acids in multiple sequence alignments of protein families can be used to fit Potts models (EV-couplings [29]) to characterize the probability of a given amino acid, given the rest of the sequence (left); see Methods for details. To compare evolutionary and H-CNN predictions for site-specific amino acid profiles, the profile overlap is computed as the centered cosine similarity between the predicted probability profiles (right); see Methods. **(F)** The profile overlaps are strongly peaked around one, implying perfect overlap in data (purple); the average profile overlap across 11,221 sites from a total of 67 protein families is 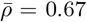. The H-CNN predictions are notably different for the shuffled data, for which the profile overlap peaks near zero (cyan), with an average of 0.002.

H-CNN predicts the conformationally unique amino acids of Glycine and Proline with over 90% accuracy. Meanwhile, amino acids with typical side-chains cluster based on their sizes and the physico-chemical properties of the side-chains including aromatic, hydrophobic, and charged groupings (Fig. 2A). The inferred amino acid preferences cluster well according to the input amino acid type (true label) in the low-dimensional UMAP representation [35], and amino acids with similar physicochemical properties cluster in nearby regions in the UMAP (Fig. 2B,C).

H-CNN predictions reflect amino acid preferences seen in evolutionary data, even though the network is not trained on multiple sequence alignments (MSAs) of protein homologs. Specifically, the interchangeability of amino acids that H-CNN predicts is 71% correlated with the substitution patterns in evolutionary data, represented by the BLOSUM62 matrix (Fig. 2D). In addition, the amino acid preferences predicted by H-CNN at each site are consistent with evolutionary preferences inferred from the covariation of residues in multiple sequence alignments of protein families [29, 36, 37]; see Fig 2E,F and Methods for details.

Ablation studies further reveal that the H-CNN’s processing of information corresponds to physical intuition. Removing information about SASA or charge from the input data results in roughly a 10% drop in classification accuracy; see Fig. S4 and the discussion on ablation studies in the Methods. Information from SASA mostly impacts the network’s ability to predict hydrophobic amino acids, with some hydrophilic amino acids (R,K,E) also impacted. When charge is removed, the network demonstrates worse predictions on charged and polar amino acids most notably R, C, N, and E.

### H-CNN learns an effective physical potential for protein micro-environments

Since H-CNN is trained to predict the most natural amino acid given its neighborhood, it should also be able to recognize an unnatural protein configuration. To test this hypothesis, we characterize the response of the H-CNN predictions to physical distortions in native atomic micro-environments. We introduce distortions through local *in silico* shear perturbation of the protein backbone at a given site *i* by an angle *δ*, resulting in a transformation of the backbone angles by *ϕ_i_* → *ϕ_i_* + *δ, ψ*_*i*–1_ → *ψ*_*i*–1_ – *δ* (Fig. 3A and Methods); this perturbation is local with minimum downstream effects [38]. We measure the distortion of the protein structure due to shear by calculating the change in the root-mean-square deviation in the pairwise distances of all atoms of the perturbed protein structure relative to that of the wild-type (RMSΔ*D_ab_*, for all pairs of atoms (*a, b*)); Fig. 3B.

**FIG. 3.**
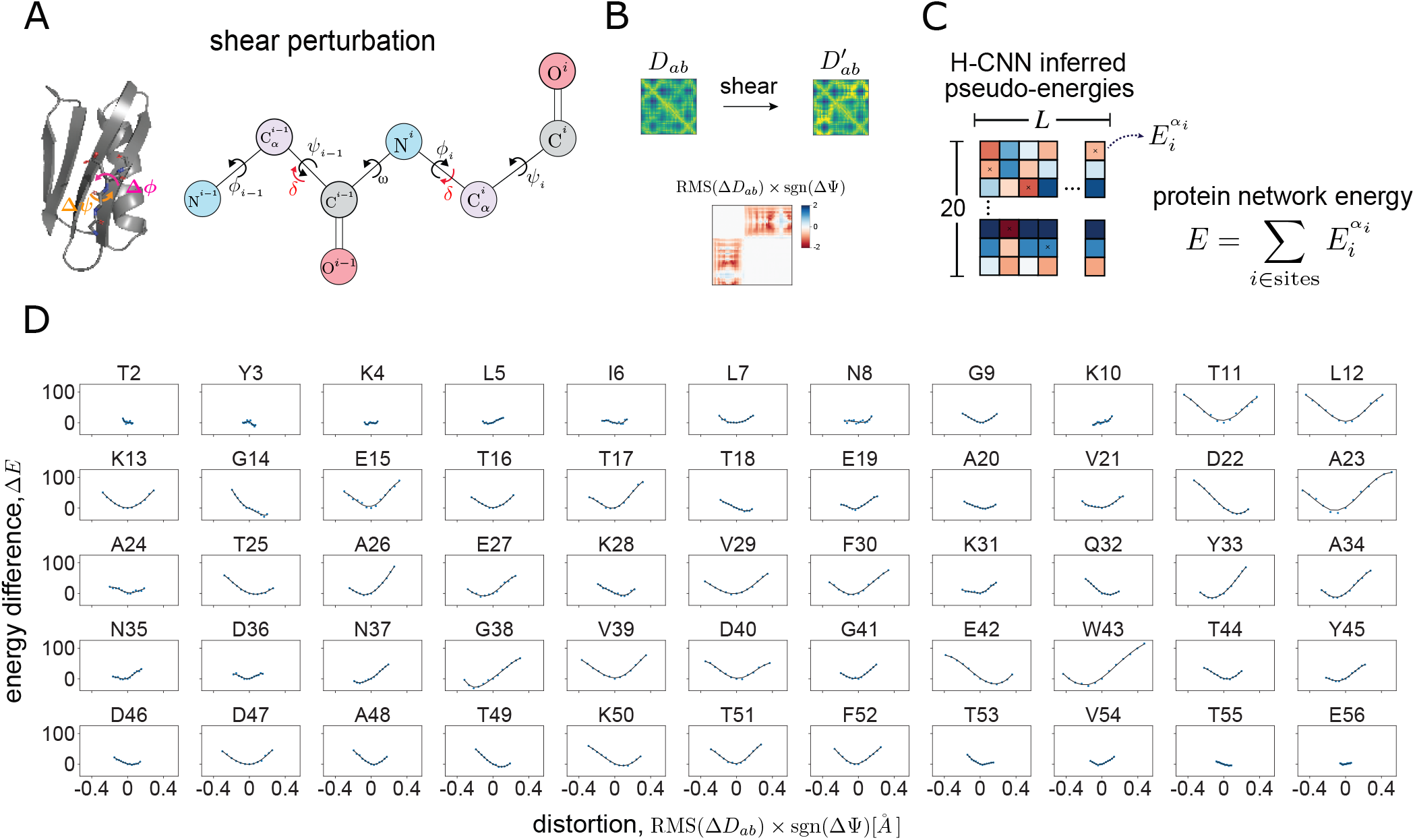
Response of H-CNN predictions to physical distortions in protein structures. **(A)** The schematic shows shear perturbation in a protein backbone by an angle *δ* at site *i* as a rotation of side-chains around the backbone by the angles [*ϕ_i_, ψ*_*i*–1_] → [*ϕ_i_* + *δ, ψ*_*i*–1_ – *δ*] [38]. **(B)** Shearing changes the pairwise distance matrix *D_ab_* between all atoms in a protein structure. The total physical distortion is computed as the root-mean-square of changes in the pairwise distances that are less than 10 Å (i.e., residues within the same neighborhood), multiplied by the sign of the change in the angle *ψ*. **(C)** For a given perturbation, the network energy *E* is determined by the sum of pseudo-energies of the wild-type amino acid at all sites in the protein, and the change in this quantity by shearing Δ*E* measures the tolerance of a structure to a given perturbation. **(D)** Panels show the change in the network energy in response to the structural distortion by shear perturbation at all sites in protein G, with the amino acid type and the site number indicated above each panel.

We measure the response of the protein to shear perturbation by analyzing the change in the logits produced by H-CNN, which we term pseudo-energies 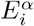 (Fig. 1). Specifically, for a given distorted structure, we re-evaluate the pseudo-energy of each amino acid in the protein, and define the total H-CNN predicted energy by summing over the pseudo-energies of all the amino acids in a protein (Fig. 3C). The change in the predicted energy of a protein due to distortion (relative to the wild-type) Δ*E* is a measure of H-CNN’s response to a given perturbation. A positive Δ*E* indicates an unfavorable change in the protein structure.

In protein G, the change in the predicted energy Δ*E* as a function of distortion in the structure RMSΔ*D_ab_* due to shearing at different sites reveals two trends (Fig. 3D). First, the protein network energy appears to respond locally quadratically to perturbations. Second, perturbations generally result in higher protein network energy, corresponding to a less favorable protein microenvironment. Taken together, by training on a classification task and by constraining the network to respect the relevant rotational symmetry, H-CNN has learned an effective physical potential for protein micro-environments in which the native structure is generally more favorable and robust to local perturbations (i.e., it is at the energy minimum).

This observation of a minimum energy extends beyond the wild-type sequence when biophysically similar amino acids are substituted in the energy sum (Methods and Fig. S3A). Notably this pattern appears not to be just an artifact of the structure since the minimum disappears when random amino acids are used to calculate the network energy (Fig. S3B).

### H-CNN predicts effect of mutations on protein stability

Characterizing amino acid preferences in a protein neighborhood is closely related to the problem of finding the impact of mutations on protein function. Here, we test the accuracy of H-CNN in predicting the stability effect of mutations in 40 different variants of the T4 lysozyme protein. Each of these variants is one amino acid away from the wild-type, with variations spanning 23 residues of the protein. Notably, the tertiary structure of the wild-type T4 lysozyme protein as well as the 40 mutants are available through different studies [30, 39–55]; see Table S3 for details on these mutants.

H-CNN predicts that the wild-type amino acids are the most favorable in the wild-type structure, while the mutant amino acids are generally more favorable in the mutant structures, regardless of their stabilizing effects (Fig. 4A). These variant-specific preferences are not surprising since the folded protein structure can relax to accommodate for amino acid changes, resulting in a structural neighborhood that is more consistent with the statistics of the micro-environments around the mutated amino acid than that of the wild-type. However, the confidence that H-CNN has in associating an amino acid with a given structural neighborhood can change depending on the stability effect of the mutation. The log-ratio of the H-CNN inferred probability for the mutant amino acid in the mutant structure versus that of the wild type amino acid in the wild type structure, Δlog *P* = log *P*_mut_/*P*_wt_ can provide an approximation to the ΔΔ*G* associated with the stability of a mutation (Methods).

**FIG. 4.**
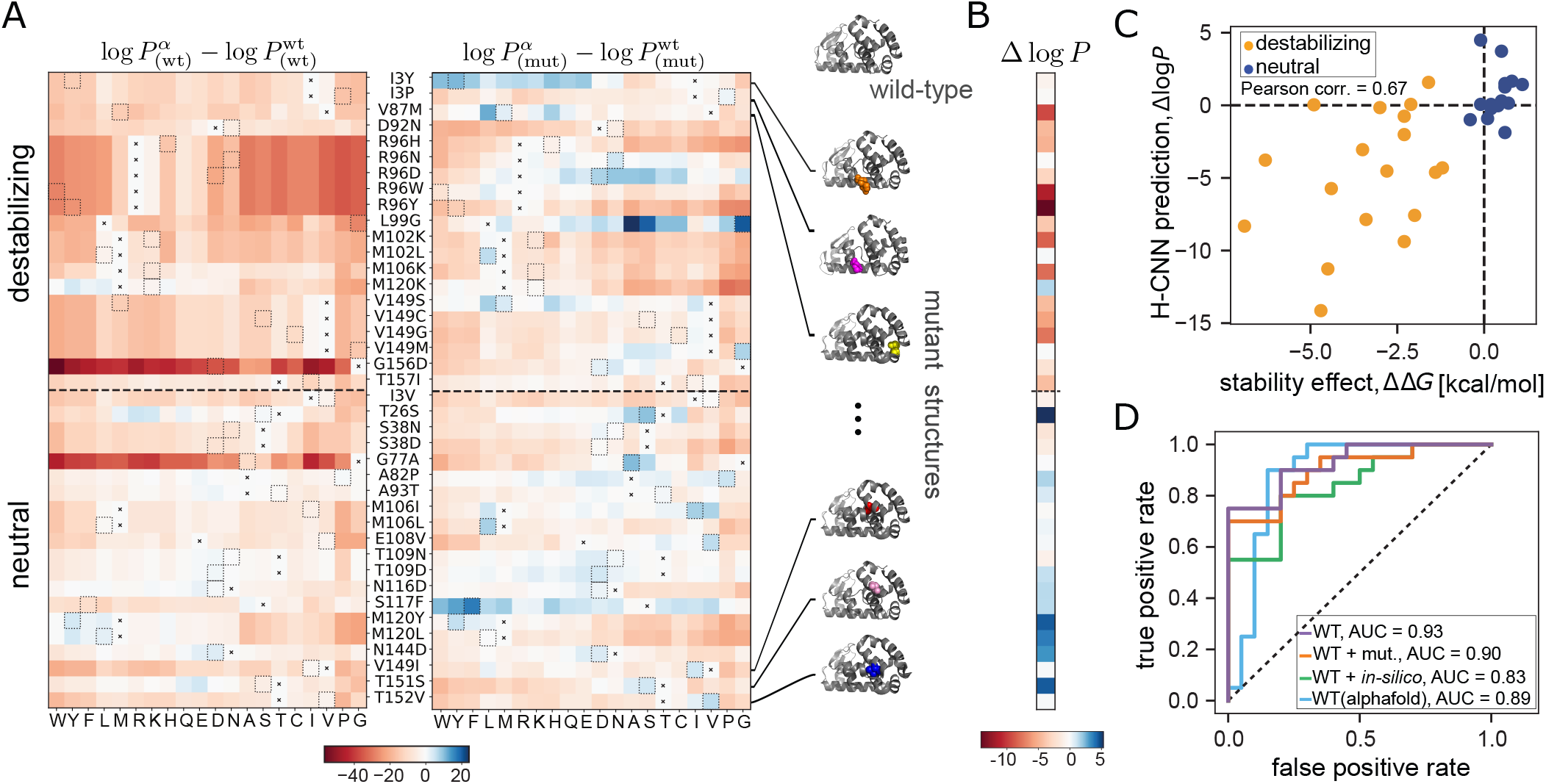
Predicting the stability effect of mutations in T4 lysozyme with H-CNN. **(A)** Heatmaps of H-CNN predicted log probability of different amino acids (columns) relative to that of the wild-type amino acid for 40 variants with single amino acid substitution from the wild-type (rows). For each variant (row), the position and the identity of the wild-type amino acid and the mutation are denoted between the two heatmaps as: wild-type, site number, mutation. The left panel shows the predictions using the wild-type protein structure (subscript (wt)), while the right panel shows the predictions using the structure of the specified mutant at each row (subscript (mut)). In each row the wild-type amino acid is indicated by an ×, and a dotted box shows the amino acid of the mutant. **(B)** The H-CNN predicted log-probability ratio 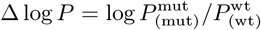 of the mutant amino acid on the mutant structure with respect to the wild-type amino acid on the wild-type structure is shown for all variants. The predicted ratios for destabilizing mutations are negative, while those for the neutral / beneficial mutations are positive. **(C)** The H-CNN predicted log-probability ratio Δlog *P* shown against the experimentally evaluated ΔΔ*G* for the stability effect of mutations in each protein structure; Pearson correlation of 67%. **(D)** The true positive vs. false positive rates (ROC curve) is shown for classification of the 40 amino acid mutations in (A) into deleterious and neutral/beneficial classes based on the H-CNN predicted log-probability ratios Δlog *P*, using the wild-type and the mutant structures (orange), only the wild-type structure (purple), the wild-type structure for the wild-type and an *in silico* relaxed structure for each mutant (green), and the AlphaFold predicted structure of the wild-type (blue). The corresponding area under the curves (AUCs) are reported in the figure.

The inferred H-CNN predicted log-probability ratio is generally negative for destabilizing mutations, and non-negative for neutral/weakly beneficial mutations (Fig. 4B). Previously, a structure-based CNN model with voxelized protein structures has shown a similar qualitative result [30]. Further quantitative analysis shows that the log-probability ratio is 67% correlated with the experimentally evaluated ΔΔ*G* values for these variants (Fig 4C). Moreover, the receiver-operating-characteristic (ROC) curves in Fig. 4D show that the log-ratio of amino acid probabilities can reliably discriminate between destabilizing and neutral mutations, with an area under the curve (AUC) of 0.90.

The availability of tertiary structures for a large number of variants is a unique feature of this dataset, and in most cases such structural resolution is not accessible. To overcome this limitation and predict the stability effect of mutations by relying on the wild-type structure alone, we used PyRosetta to relax the wild-type T4 lysozyme structure around a specified amino acid change [38] (Methods). We find that the log-probability ratios 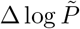 estimated based on these *in silico* relaxed mutant structures are mostly negative (non-negative) for destabilizing (neutral) mutations (Fig. S5) and are correlated with the stability effect of mutations ΔΔ*G* (Fig. S5). However, structural relaxation can add noise to the data, causing the protein micro-environments to deviate from the natural structures that H-CNN is trained on. Thus, using the *in silico* relaxed structures slightly reduces the discrimination power of our model between deleterious and near-neutral mutations (AUC = 0.83); see Fig. 4D.

In contrast, the preferences estimated based on the wild-type structure only can discriminate between destabilizing and neutral mutations very well, even though most mutations are inferred to be deleterious with respect to the wild-type (AUC = 0.93 in Figs. 4D, S5). In other words, by using the wild-type structure only, our model can predict the relative stability effect of mutations correctly but not the sign of ΔΔ*G* (Fig. S5). Indeed, our inferred log-probability ratios based on the wild-type structure show a substantial correlation of 64% (Pearson correlation) with the stability effect of a much larger set of 310 single point mutants [56], for which protein structures are not available (Fig. S6).

When no experimentally determined structure is available, computationally resolved protein structures from AlphaFold can also be used to predict the stability effect of mutations. The H-CNN predictions using the template-free AlphaFold2 predicted structure of T4 lysozyme wild-type sequence display substantial discrimination ability between destabilizing and near-neutral mutations (Fig. 5) and are correlated with the mutants’ ΔΔ*G* values (Figs. S5, S6).

**FIG. 5.**
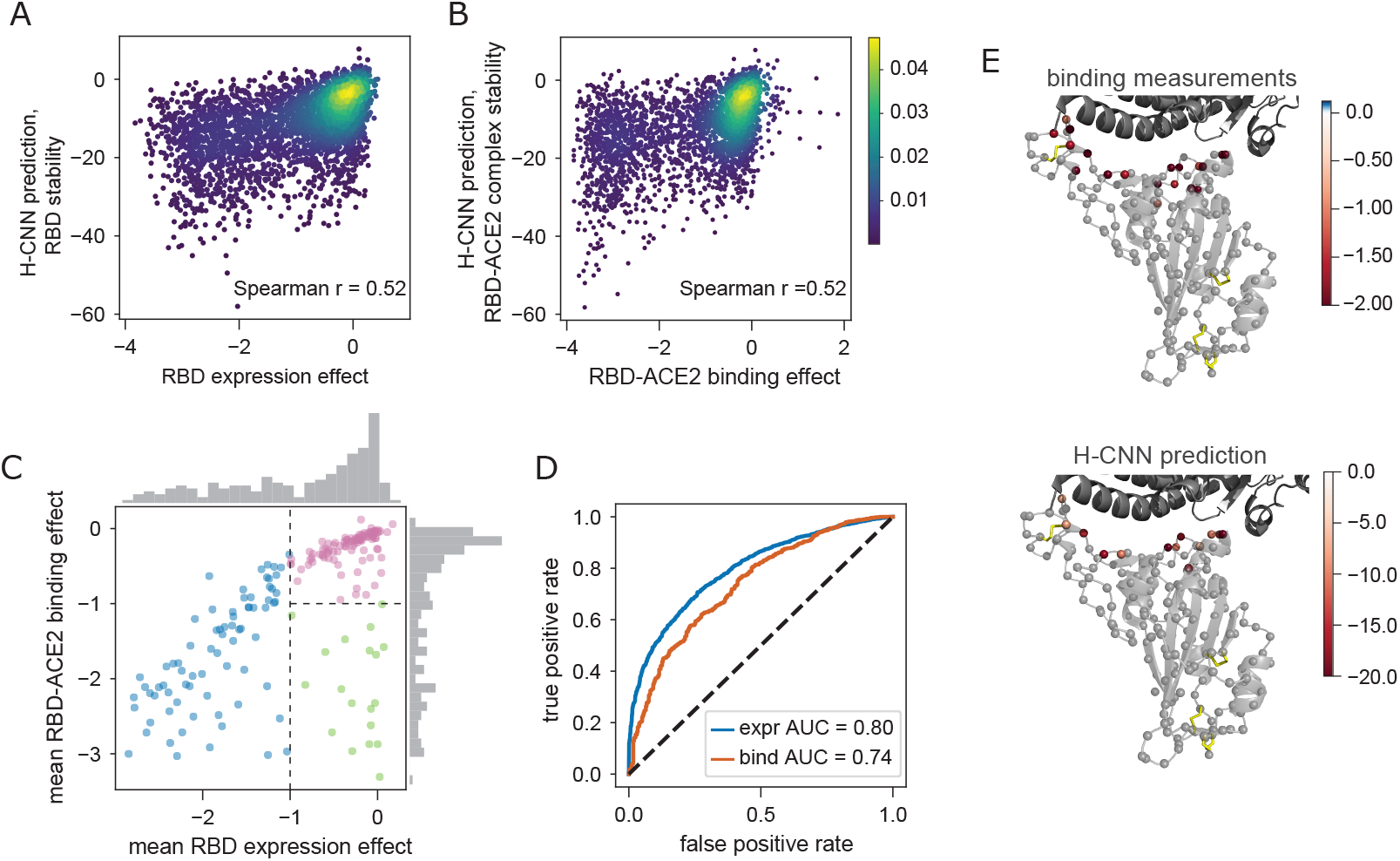
Predicting the stability and binding of the RBD protein of SARS-CoV-2 with H-CNN. **(A)** The density plot shows H-CNN predictions for the RBD stability, using the isolated protein structure of RBD, against the mutational effects on the RBD expression from the DMS experiments; Spearman correlation *r* = 0.52. **(B)** The density plot shows H-CNN predictions for the RBD binding to the ACE2 receptor, using the co-crystallized RBD-ACE2 protein structure, against the DMS measurements for mutational effects on binding; shared color bar for (A) and (B). **(C)** The mean effect of mutations at each site on the RBD-ACE2 binding is shown against the mean effect on the RBD expression. The histograms show the corresponding distribution of effects across sites along each axis. The categories are shown: (i) sites that are intolerant to mutations due to destabilizing effect, i.e., low expression (blue), (ii) sites that are tolerant of mutations for expression but not binding (green), and (iii) sites that are tolerant of mutations for both expression and binding (pink). **(D)** Blue: true positive vs. false positive rate (ROC curve) for classification of amino acid mutations into stable (expr > –1) vs. unstable (expr < –1), based on the H-CNN predictions using the isolated RBD structure; AUC =0.8. Red: the ROC curve for mutation classification into bound (bind > –1) vs. unbound (bind < –1), based on the H-CNN predictions using the co-crystallized RBD-ACE2 structure; AUC= 0.74. **(E)** The effect of mutations on binding from the DMS experimental data for the green sites in (C) (top) and the corresponding H-CNN predictions from the RBD-ACE2 structure complex for sites identified by H-CNN in Fig. S10 to be tolerant of mutations for stability but not binding (bottom) are shown throughout the structure.

### H-CNN predicts fitness effect of mutations for binding of SARS-CoV2 to the ACE2 receptor

Recent deep mutational scanning (DMS) experiments measured the effect of thousands of mutations in the receptor-binding domain (RBD) of SARS-CoV-2 on the folding of the RBD (through expression measurements) and its binding to the human Angiotensin-Converting Enzyme 2 (ACE2) receptor [57, 58].

H-CNN can be used to predict the effect of mutations on RBD, either in isolation or bound to the ACE2 receptor. The former can be interpreted as the effect of mutations on the stability of RBD, which is measured by the expression of the folded domain in the experiments [57–59], while the latter can be used to characterize amino acid preferences for binding at the RBD-ACE2 interface. Fig. 5A,B shows that the H-CNN predictions are correlated with the stability and binding measurements in the DMS experiments from ref. [58]; site specific effects are depicted in Figs. S8, S9.

The average effect of mutations on expression and binding can define three categories of sites and/or mutations (Fig. 5C): (i) sites that are intolerant to mutations (due to destabilizing effects) and show a substantially reduced expression of mutants (blue), (ii) sites that are tolerant of mutations for expression but not binding (green), and (iii) sites that are tolerant of mutations for both expression and binding (pink). Using the isolated structure of RBD, H-CNN can well classify mutations according to their stability effect (AUC = 0.8; Fig. 5D). Similarly, with the structure of the RBD-ACE2 complex, H-CNN can classify mutations according to their tolerance for binding (AUC = 0.74; Fig. 5D).

Expectantly, the sites that are tolerant of mutations for expression but not binding (green category from the DMS data in Fig. 5C) are located at the interface of the RBD-ACE2 complex, and H-CNN correctly predicts this composition (Fig. 5E, Fig. S10). The overall impact of mutations on binding for these sites is shown in Fig. 5E.

Identifying candidate sites that can tolerate mutations and can potentially improve binding is important for designing targeted mutagenesis experiments. Instead of agnostically scanning single point and (a few) double mutations over all sites, these predictions can inform experiments to preferentially scan combinations of viable mutations on a smaller set of candidate sites. In previous work, evolutionary information was used to design such targeted mutagenesis for the HA and NA proteins of influenza [60, 61]. A principled structure-based model could substantially improve the design of these experiments.

## Discussion

The success of AlphaFold has demonstrated the power of machine learning in predicting protein structure from sequence [3]. The challenge now is to leverage the experimentally and computationally determined protein structures to better understand and predict protein function. Our H-CNN model is a computationally powerful method to represent protein tertiary structures, and characterizes local biophysical interactions in protein micro-environments. Our model is physically motivated in that it respects rotational symmetry of protein structure data, allowing for significantly faster training time compared to previous approaches [30, 31].

Similar to recent language models, H-CNN also demonstrates strong cross-task generalization by predicting quantitative effects of amino acid substitutions on function (i.e., zero-shot predictions), including protein stability or binding of protein complexes. Generally, massive language models trained on large and diverse protein sequence databases are shown to generalize well to predict mutational effects in proteins without any supervision [6, 7, 10, 62, 63]. State-of-the-art methods include ESM-1b for zero-shot predictions [10] and MSA transformers that use evolutionary information from MSAs of protein families to predict the effect of mutations [62]. The benchmark for these methods is the large set of DMS experiments, for which most zero-shot sequencebased predictions show an average accuracy of about 50% in predicting the rank order of the mutational effects [63]. Our structure-based H-CNN method shows a comparable accuracy in predicting the mutational effect in DMS experiments of the RBD protein in SARS-CoV-2, yet with much fewer parameters; a more systematic analysis would be necessary to compare these different approaches. Nonetheless, it would be interesting to see how the features extracted by H-CNN can complement the sequence-based language models to potentially improve zero-shot predictions for mutational effects in proteins.

Recent work has shown that combining structural data with evolutionary information from MSAs in deep learning models can be powerful in predicting mutational effects in proteins [64]. We have shown that H-CNN recapitulates the functional information reflected in evolutionary data, further reinforcing the idea that physically guided structure-based machine learning models could be sufficient in predicting protein function, without a need for MSAs. Importantly, our MSA-independent approach enables us to apply H-CNN to protein structures with no available homologs, including the *de novo* protein structures.

The H-CNN learned representations of amino acid neighborhoods could be used as input to a supervised algorithm to learn a more accurate model for mutational effects in proteins; a similar approach has been used to model the stability effect of mutations in ref. [65]. Moreover, the all-atom representation of protein structures used to train H-CNN allows for generalizability, e.g. using the inferred model to analyze non-amino acid molecules or extending the model and accommodate other elements to study protein-drug or protein-DNA interactions.

Solving the inverse protein folding problem by designing a sequence that folds into a desired structure is a key step in protein design. Recent deep learning methods, including protein MPNN [16] and transformer-based ESM-IF1 [12, 15], have shown promise in designing viable sequences with a desired fold for *de novo* proteins. H-CNN’s ability to learn an effective potential in protein micro-environments merits investigation as to whether similar techniques can be used to solve the inverse folding problem for *de novo* proteins.

The learned representation of protein microenvironments with H-CNN enables us to characterize the preferences of different amino acid compositions in a structural neighborhood. Additionally, these rotationally equivariant representations could be used as building blocks of larger protein structure units, e.g. to characterize how different molecular features on a protein surface could determine its interactions with other proteins. A study in this direction could shed light on the structure-to-function map of the protein universe.

## II. ACKNOWLEDGMENT

This work has been supported by the National Institutes of Health MIRA award (R35 GM142795), the CAREER award from the National Science Foundation (grant No: 2045054), the Royalty Research Fund from the University of Washington (no. A153352), the Microsoft Azure award from the eScience institute at the University of Washington. This work is also supported, in part, through the Department of Physics, and the College of Arts and Sciences at the University of Washington.

