## Supplementary Material for "Learning the shape of protein micro-environments with a holographic convolutional neural network"

CONTENTS

I. Data pre-processing and code availability 1

II. Rotationally equivariant structure-to-function map for proteins 2
A. Mathematical foundation for 3D rotationally equivariant transformations 2
B. Rotationally equivariant encoding of protein micro-environments as spherical holograms 5
C. Rotationally equivariant model of protein micro-environments with H-CNN 5

III. Network and training procedure for H-CNN 7

IV. Comparing H-CNN with evolutionary models for amino acid preferences 8

V. Effect of shear perturbation on protein micro-environments 8

References 9

### I. DATA PRE-PROCESSING AND CODE AVAILABILITY

**Code and data availability.** All codes and references to data are available on github through: https://github.com/StatPhysBio/protein_holography.

**Processing of the protein structure data.** We minimally process the protein structures from the Protein Data Bank (PDB) [1]. We use PyRosetta [2] to infer coordinates of hydrogen atoms in the protein structure. We also use PyRosetta to assign solvent accessible surface area (SASA) and partial charge to all atoms.

**Constructing the training, the validation, and the test sets for protein neighborhood classifications.** We classified amino acid neighborhoods extracted from the tertiary protein structures from the PDB. We used ProteinNet's splittings for training and validation sets of the CASP12 competition to avoid redundancy due to similarities in homologous protein domains [3]. Since PDB identifiers in ProteinNet were only provided for the training and the validation sets, we used ProteinNet's training set as the base of both our training and validation and ProteinNet's validation set as our testing set. Specifically, to define our training and validation sets, we make a [80%, 20%] split in the ProteinNet's training data of proteins with X-ray crystallography structures of 2.5Å resolution or better. Furthermore, in anticipation of validating our model on experimental stabilities of T4 lysozyme and SARS-CoV2 receptor-binding domain (RBD), we removed all structures that belonged to the same UniProt family as the domains from each of these proteins. We used all of ProteinNet's validation structures as our testing set.

This splitting resulted in 10,957 training structures, 2,730 validation structures, and 212 testing structures, each of which contained 2,810,503, 682,689, and 4,472 neighborhoods, respectively.

**Data processing to assess the stability effect of mutations in the T4 lysozyme protein.** Predictions for the stability effect of mutations in the T4 lysozyme protein were made using the PDB structure 2LZM [4] as the wild-type. The PDB identifiers for structures of all the T4 lysozyme variants used in our analyses (shown in Figs. 4 and S6) along with the reported $\Delta\Delta G$ and the pH values of the stability measurements are reported in Table S3.

Experimental measurements of $\Delta\Delta G$ values in variants for which we do not have a matching protein structure (shown in Fig. S7) are taken from the dataset FireProtDB [5].

The AlphaFold prediction of the T4 lysozyme protein structure, used in Fig. 4C and S7, was performed using the AlphaFold GoogleColab notebook [6].

Relaxation of mutant T4 lysozyme structures for the *in silico* model in Figs. 4B,C and S6, was done using PyRosetta `FastRelax` with the score function `ref2015_cart` [2]. Backbone flexible positions were restricted to the mutated residue and the neighboring residues in the sequence. Sidechain flexible positions were restricted to the neighbors with alpha carbons within 10 Å of the mutated residue's alpha carbon.

**Data processing to assess the fitness effect of mutations in the RBD of SARS-CoV-2.** Deep mutational scanning experiments from ref. [7] were used to obtain the SARS-CoV-2 RBD expression and binding to the ACE2 receptor. Predictions of the RBD-ACE2 binding strength, shown in Fig. 5B, were made using the co-crystallized protein structure with the PDB identifier 6M0J [8]. The predictions for the RBD stability, shown in Fig. 5A, were made using the 6M0J structure with the ACE2 chain computationally removed. This computational removal was performed by selecting the RBD structure in PyMol [9] and exporting a PDB file from said selection.

**Data processing to assess the effect of shearing in Protein G.** To analyze the effect of shearing in protein G, shown in Figs. 3 and S5, we used the native structure of protein G with the PDB identifier 1PGA [10]. Sheared structures were produced with PyRosetta by manually editing the backbone angles $\phi$ and $\psi$ of each residue (Fig. 3A) via `Pose.set_phi` and `Pose.set_psi` [2].

### II. ROTATIONALLY EQUIVARIANT STRUCTURE-TO-FUNCTION MAP FOR PROTEINS

The tertiary protein structure is a result of a folding process whereby amino acid sidechains interact with each other and the backbone structure to form the three-dimensional (3D) shape of the protein. The favorability of a certain amino acid occurring at a given site in a protein is determined by numerous factors: the presence of this amino acid during translation, the ability of the polypeptide chain to fold with that amino acid present, the amino acid's interactions with the local structural surroundings of the protein, and the amino acid's interactions with the surrounding environment. Our work focuses on revealing the third factor's contribution. Specifically, we aim to learn how a micro-environment in a protein's tertiary structure (i.e., atoms in a given structural neighborhood) constrain the usage of different amino acids in the neighborhood.

We define a structural neighborhood surrounding a focal amino acid by the coordinates (and identities) of all the $N$ atoms from the surrounding amino acids in the structure, within a given radius of the focal residue. Mathematically, we want to find a map $f : \mathbb{R}^{N\times 3} \to [\mathbb{R}^+_{\leq 1}]^{19}$ between the 3D coordinates of the $N$ neighboring atoms and the vector of probabilities $p \in [\mathbb{R}^+_{\leq 1}]^{19}$ associated with preferences for different types of amino acids that can be placed at the center of the neighborhood; note that the amino acid probability vector has 19 independent entries due to normalization.

Assuming that the protein structure is a crystal (at least on average), the local interactions between the amino acid at the focal site of interest and the surrounding ones in the protein structure have rotational symmetry, in that they only depend on the pairwise distances of atoms and not on their absolute orientation in space. This symmetry implies that the interactions are unchanged under global rotations since a global rotation preserves pairwise distances and relative orientations. Therefore, any model of amino acid preferences based on local structural information should respect 3D rotational symmetry. Thus, our goal becomes learning a map $f$, as described above, that is also invariant to global rotations acting on the inputs. A rotation acting on any Cartesian vector can be expressed as an Euler angle matrix $R_{\text{euler}}$ (Fig. S1). Since our inputs $\mathbf{x}$ for coordinates of the atoms in a structural neighborhood are Cartesian vectors $\mathbf{x} \in \mathcal{R}^{N\times 3}$, a global rotation $R$ would act on these vectors via a Euler angle matrix, which we represent by $D_{\text{cart.}}(R)$. Therefore, the rotationally invariant map should have the property $f(D_{\text{cart.}}(R)\mathbf{x}) = f(\mathbf{x})$.

In the rest of this section, we describe the mathematical foundation to construct 3D rotationally equivariant models for protein structure data. In the following section (Section II A), we specifically focus on our approach in encoding the structural data and training neural networks to learn 3D rotationally equivariant models for protein neighborhoods. The methodology discussed in Sections II B, II C is self-contained, and if a reader wishes they can choose to skip Section II A on the mathematical foundation of our approach.

#### A. Mathematical foundation for 3D rotationally equivariant transformations

The simplest approach to train a neural network model that is invariant to global rotations of proteins is to use invariant descriptors of the tertiary structure as inputs to the neural network. For example, one can use dot products between all Cartesian vectors describing the coordinates of atoms within a structural neighborhood. However, recent work has shown that neural networks that are allowed to work with full geometric data as opposed to simple invariants have higher expressivity and perform better on learning tasks [11].

Networks that build more complex invariants rely on representation theory and the idea of *equivariance* to build more expressive models. Equivariance is a generalization of invariance. Rather than requiring no change when inputs are transformed, equivariance allows the outputs of a map to change but requires them to change in a well-defined way. Under a transformation of the inputs corresponding to a rotation $R$, an equivariant map $f$ satisfies the relation

$$f(D_{\text{input}}(R)x) = D_{\text{out}}(R)f(x) \tag{S1}$$

where $D_{\text{input}}(R)$ is the operator corresponding to the rotation in the input space, and $D_{\text{out}}(R)$ is the operator corresponding to the rotation in the output space. Note that if $D_{\text{out}}(R)$ is the identity matrix for all rotations, then the map is invariant.

The key to building an equivariant map is to know the operator $D_{\text{out}}$ for the transformation of interest. The prevailing method in the field of equivariant machine learning is to modularly build a neural network out of equivariant units where each unit has an input and output space where these operators are known. The input and output space are often the same and tend to be the Fourier space defined by the symmetry that is respected.

Rotations in three dimensions are non-commutative which means that the output of two consecutive rotations can depend on the order by which they act on the input vector. As a result, the Fourier space for rotations is more complicated thaN that of a commutative transformation like translations. Here, we will first use translations as an illustrative example for equivariant operations in Fourier space and then generalize to rotations.

**Translational equivariance.** The Fourier transform of a signal on a real line $f(x)$ can be expressed as $\hat{F}(k) = \int_{-\infty}^{\infty} e^{-ikx} f(x)dx$. We define a translated signal $f_a(x) = f(x-a)$ which is the original signal translated by a distance $a$. The Fourier transform of this translated signal follows

$$\begin{aligned}\hat{F}_a(k) &= \int_{-\infty}^{\infty} e^{-ikx} f_a(x)dx \\ &= \int_{-\infty}^{\infty} e^{-ikx} f(x-a)dx \\ &= \int_{-\infty}^{\infty} e^{-ik(x'+a)} f(x')dx' \\ &= e^{-ika}\hat{F}(k)\end{aligned} \tag{S2}$$

Thus, the Fourier transform of a translated signal transforms as $\hat{F}_a \xrightarrow{a} e^{-ika}\hat{F}(k)$, implying that the Fourier transform is equivariant under translation with an output operator that is simply a phase shift $D_{\text{out}}(a) = e^{-ika}$. These operators $D_{\text{out}}(a)$ form a representation of the group of 1D translations [12].

In three dimensions, translations act on vectors via $\mathbf{x} \to \mathbf{x} + \mathbf{a}$ and the representation is given by the operators operator $D_{\text{out}}(\mathbf{a}) = e^{-i\mathbf{k}\cdot\mathbf{a}}$.

**Rotational equivariance.** Similar to translation, we can use Fourier space to define an equivariant transformation for rotations. We consider rotations about a given reference point that defines an origin for a spherical coordinate system in which points in 3D can be expressed by the resulting spherical coordinates $(r, \theta, \phi)$. Since the radius $r$ (i.e., the distance of a point from to the reference) does not change under rotations about the origin, we will ignore the radial component for now and consider a signal over the sphere of radius $r$, $f(\theta, \phi) : \; S^2(r) \to \mathbb{R}$. The Fourier transform $\hat{F}$ of the signal on the sphere follows,

$$\hat{F}_{\ell m} = \int_0^{2\pi} \int_0^{\pi} f(\theta,\phi) Y_{\ell m}(\theta,\phi) \sin\theta \, \mathrm{d}\theta \, \mathrm{d}\phi \tag{S3}$$

where $Y_{\ell m}(\theta, \phi)$ is the spherical harmonic of degree $\ell$ and order $m$ defined as

$$Y_{\ell m}(\theta,\phi) = \sqrt{\frac{2n+1}{4\pi}\frac{(n-m)!}{(n+m)!}} e^{im\phi} P_\ell^m(\cos\theta) \tag{S4}$$

where $\ell$ is a non-negative integer ($0 \leq \ell$), and $m$ is an integer within the interval $-\ell \leq m \leq \ell$. $P_\ell^m(\cos\theta)$ is the Legendre polynomial of degree $\ell$ and order $m$, which, together with the complex exponential $e^{im\phi}$, define sinusoidal functions over the angles $\theta$ and $\phi$. In quantum mechanics, spherical harmonics are used to represent the orbital angular momenta, e.g. for an electron in a hydrogen atom. In this context, the degree $\ell$ relates to the eigenvalue of the square of the angular momentum, and the order $m$ is the eigenvalue of the angular momentum about the azimuthal axis.

The operators that describe how spherical harmonics transform under rotations are called the Wigner D-matrices, denoted by $D^{\ell}_{mm'}(R)$ [12]; this operator is a matrix because the rotation group in 3D is not commutative. Under a rotation $R$, spherical harmonics transform as

$$Y_{\ell m}(\theta,\phi) \xrightarrow{R} \sum_{m'=-\ell}^{\ell} D^{\ell}_{m'm}(R) Y_{\ell m'}(\theta,\phi) \tag{S5}$$

Since spherical harmonics transform in this well-defined way, we can now examine how a rotation acts on the Fourier transform of a signal over the sphere. Suppose we have a signal $f(\mathbf{n})$ defined on the elements $\mathbf{n}$ of the spherical shell $S^2$ (i.e., the set of angular coordinates for points on the sphere). The Fourier coefficients associated with the rotated signal $f_R(\mathbf{n}) = f(R\mathbf{n})$ follow,

$$\begin{aligned}
\hat{F}^{R}_{\ell m} = \int Y_{\ell m}(\mathbf{n}) f_R(\mathbf{n}) \mathrm{d}\Omega &= \int Y_{\ell m}(\mathbf{n}) f(R\mathbf{n}) \mathrm{d}\Omega \\
&= \int Y_{\ell m}(-R\mathbf{n}') f(\mathbf{n}') \mathrm{d}\Omega' \\
&= \int \sum_{m'=-\ell}^{\ell} D^{\ell}_{m'm}(-R) Y_{\ell m}(\mathbf{n}') f(\mathbf{n}') \mathrm{d}\Omega' \\
&= \sum_{m'=-\ell}^{\ell} D^{\ell}_{m'm}(-R) \int Y_{\ell m}(\mathbf{n}') f(\mathbf{n}') \mathrm{d}\Omega' \\
&= \sum_{m'=-\ell}^{\ell} D^{\ell}_{m'm}(-R) \hat{F}_{lm}
\end{aligned} \tag{S6}$$

where $\mathrm{d}\Omega = \sin^2\theta \, \mathrm{d}\theta \, \mathrm{d}\phi$ is the angular differential in the spherical coordinate system. Under a rotation $R$ about the origin, the spherical Fourier coefficients of a real-space signal transform according to $\hat{F}_{\ell m} \xrightarrow{R} \sum_{m'=-\ell}^{\ell} D^{\ell}_{m'm}(-R) \hat{F}_{\ell m'}$, which is simply a matrix product.

It should be noted that the Wigner D-matrices form an irreducible representation of SO(3), which is the group of all rotations around a fixed origin in three-dimensional Euclidean space. Therefore, it provides a natural basis to expand functions in SO(3) while respecting the rotational symmetry.

**Nonlinear transforms respecting rotational equivariance.** One key feature of neural networks is applying nonlinear activations, which enable a network to approximately model complex and non-linear phenomena. Commonly used nonlinearities include reLU, tanh, and softmax functions. However, these conventional non-linearities can break rotational equivariance in the Fourier space. To construct expressive rotationally equivariant neural networks we can use the Clebsch-Gordan tensor product, which is the natural nonlinear (or bilinear in the case of using two sets of Fourier coefficients) operation in the space of spherical harmonics [12].

Given two spherical tensors $\hat{F}_{\ell_1}$ and $\hat{G}_{\ell_2}$, we can take a product between all of their components $\hat{F}_{\ell_1 m_1}\hat{G}_{\ell_2 m_2}$, which would allow us to express nonlinearities. Although these products do not behave well under rotations, linear combinations of them do transform in well-defined ways. The Clebsch-Gordan coefficients are the coefficients that decompose these products back into the space of spherical harmonics, where Wigner D-matrices define the rotation operations. Specifically, the product of spherical tensors $\hat{F}_{\ell_1}$ and $\hat{G}_{\ell_2}$ will yield spherical tensors of degree $L$ such that $|\ell_1 - \ell_2| < L < \ell_1 + \ell_2$ in the following way,

$$\hat{H}_{LM} = \sum_{m_1=-\ell_1}^{\ell_1} \sum_{m_2=-\ell_2}^{\ell_2} \langle LM | \ell_1 m_1; \ell_2 m_2 \rangle \hat{F}_{\ell_1 m_1} \hat{G}_{\ell_2 m_2} \tag{S7}$$

where $\langle LM | \ell_1 m_1; \ell_2 m_2 \rangle$ is a Clebsch-Gordan coefficient and can be precomputed for all degrees of spherical tensors [12]. Similar to spherical harmonics, Clebsch-Gordan products also appear in quantum mechanics, and they are used to express couplings between angular momenta. In following with recent work on group-equivariant machine learning [11, 13–15], we will use Clebsch-Gordan products to express nonlinearities in 3D rotationally equivariant neural networks for protein structures.

#### B. Rotationally equivariant encoding of protein micro-environments as spherical holograms

We define an amino acid's micro-environment as all the atoms associated with the surrounding amino acids within a 10Å of the central residue's $\alpha$-carbon. We use the position of the central residue's $\alpha$-carbon as the reference point of the neighborhood $\mathcal{N}$. Each atom $i$ within this neighborhood, located at position $\mathbf{r}_i$ with respect to the reference point, has three attributes associated with it: (i) the element type of the atom, $e_i \in \{C, N, O, S, H\}$, (ii) the partial charge of the atom $Q_i$, and (iii) the SASA $A_i$ of the atom. With these characteristics we define a feature vector $v_i^c$ where $c$ indexes the input channels $\{C, N, O, S, H, \text{charge}, \text{SASA}\}$. This vector takes on values given by

$$v_i^c = \begin{cases} \delta_{e_i,c} & \text{for } c \in \{C, N, O, S, H\} \\ Q_i & \text{for } c = \text{charge} \\ A_i & \text{for } c = \text{SASA} \end{cases}$$

where $\delta_{ij}$ is a Kronecker delta function, and takes the value 1 if $i = j$ and 0 otherwise. Thus, for $c \in \{C, N, O, S, H\}$ the feature vector $v_i^c$ takes value 1 if atom $i$ is of type $c$ and 0 otherwise. $Q_i$ and $A_i$ denote the partial charge and the SASA (in units of $\text{Å}^2$) of atom $i$, respectively. These quantities are computed by PyRosetta for each protein structure [2].

Each channel defines a point cloud with the attributed values stored at the coordinates of the corresponding atoms. We express the point cloud of each channel in a spherical coordinate system with the origin set at the alpha carbon of the central amino acid in the neighborhood. We then define an atomic density as a sum of Dirac delta distributions parameterized by each atomic coordinate in the neighborhood $\rho^c(\mathbf{r}) = \sum_{i\in\mathcal{N}} v_i^c \delta^{(3)}(\mathbf{r} - \mathbf{r}_i)$. Here $\delta^{(3)}(\mathbf{x})$ denotes a normalized Dirac delta probability density in 3D, which is zero everywhere except at the origin $\mathbf{x} = \mathbf{0}$, where it is infinite. We use 3D Zernike transforms to encode the point clouds associated with each channel in a 3D rotationally equivariant fashion in the spherical Fourier space:

$$\hat{Z}^c_{n\ell m} = \int \rho^c(\mathbf{r}) Y_{\ell m}(\theta, \phi) R_{n\ell}(\mathbf{r})\, \mathrm{d}\Omega. \tag{S8}$$

where $\mathrm{d}\Omega$ is the angular differential in the spherical coordinate system, $Y_{\ell m}(\theta, \phi)$ is the spherical harmonics (eq. S4), and $R_{n\ell}(\mathbf{r})$ is the radial function of the 3D Zernike transform, which can be expressed as,

$$R_{n\ell}(\mathbf{r}) = (-1)^{\frac{n-\ell}{2}} \sqrt{2n+3} \binom{\frac{n+\ell+3}{2} - 1}{\frac{n-\ell}{2}} |\mathbf{r}|^\ell \, {}_2F_1\left(-\frac{n-1}{2}, \frac{n+\ell+3}{2}; \ell + \frac{3}{2}; |\mathbf{r}|^2\right) \tag{S9}$$

where ${}_2F_1(\cdot)$ is the ordinary hypergeometric function. The index $n \geq 0$ is a non-negative integer, and the radial function $R_{n\ell}(\mathbf{r})$ is nonzero only for even values of $n - \ell \geq 0$. Zernike polynomials form a complete orthonormal basis in 3D and, therefore, can be used to expand and retrieve any 3D shape, if large enough $\ell$ and $n$ coefficients are used. Approximations that restrict the series to finite $n$ and $\ell$ are often sufficient for shape retrieval, and hence, desirable algorithmically. Importantly, expansion of an input signal by Zernike polynomials (eq. S8) is rotationally equivariant since a rotation $R$ transforms the resulting coefficients through Wigner-D matrices as $\hat{Z}^c_{n\ell m} \xrightarrow{R} \sum_{m'=-\ell}^{\ell} D^\ell_{m'm}(-R)\hat{Z}^c_{n\ell m'}$ (eq. S5).

Zernike projections in the spherical Fourier space can be thought of as a superposition of spherical holograms of an input point cloud, and thus, we term this operation the *holographic encoding* of protein micro-environments.

#### C. Rotationally equivariant model of protein micro-environments with H-CNN

We use the coefficients of the Zernike expansion $\hat{Z}^c_{nlm}$ in eq. S8 (i.e., the holographic encoding of the data) as inputs to a rotationally equivariant convolutional neural network to classify amino acids based on their surrounding atomic micro-environments. We term this network holographic convolutional neural network (H-CNN).

The convolutional neural network that we use for our analysis is similar to the Clebsch-Gordan network (CGNet), described in ref. [13]. This network is built out of three rotationally equivariant modular units: (i) linear transformation of spherical harmonics, (ii) Clebsch-Gordan nonlinearity, and (iii) spherical batch normalization.

Below we will describe the computations done in each of these modular equivariant units. Without a loss of generality, we denote the operations in each unit of the network in terms of a general equivariant signal $\hat{F}^k_{lm}$ where $k$ is a catch-all index that represents all non-rotational indices (i.e., channel $c$ and index $n$ in eq. S8), and $\ell$ and $m$ correspond to the angular indices (eq. S4). The output of each unit will be denoted by $\hat{G}^k_{lm}$.

**Linear transformations in CG-nets.** Linearities are ubiquitous in neural networks and contribute partially to their powerful expressiveness. Our linearity takes the form

$$\hat{G}^{k}_{\ell m} = \sum_{k'} W^{(\ell)}_{kk'} \hat{F}^{k'}_{\ell m} \tag{S10}$$

where $W^{(\ell)}_{kk'}$ denotes two dimensional matrices that mix different input channels $k'$ (e.g. combinations of radial index $n$ and channel $c$ in the input or general learned representations in intermediate layers) producing different output channels $k$. To preserve rotational symmetry, we will impose the restriction of only combining information (i.e., forming linear combinations of inputs) that belongs to the same irreducible representations (irreps) of SO(3) in eq. S10. In other words, we mix only inputs associated with the same degree $\ell$ and order $m$ of spherical harmonics. These inputs are mixed across channels associated with different chemical or radial information in the first layer, or composite channels in the following layers. Generally, different weights are learned for each irrep $\ell$ but shared across coefficients of different order $m$.

**Clebsch-Gordan nonlinearity.** The last equivariant component of the network is a Clebsch-Gordan nonlinearity. Nonlinearities are ubiquitous in neural networks. The Clebsch-Gordan nonlinearity is the simplest equivariance-respecting nonlinearity, and it is expressive enough for most learning tasks [13, 15]. This Clebsch-Gordan nonlinearity is simply a product of spherical tensors that is decomposed back into the spherical harmonic basis via the Clebsch-Gordan coefficients,

$$\hat{H}^{K}_{LM} = \sum_{m_1=-\ell_1}^{\ell_1} \sum_{m_2=-\ell_2}^{\ell_2} \langle LM|\ell_1 m_1; \ell_2 m_2\rangle \hat{F}^{k_1}_{\ell_1 m_1} \hat{G}^{k_2}_{\ell_2 m_2} \tag{S11}$$

where $\langle LM|\ell_1 m_1; \ell_2 m_2\rangle$ is a Clebsch-Gordan coefficient and $K = (k_1, k_2)$ is the new channel index.

Generally we have a choice for which combinations of values $k_1, k_2, \ell_1$, and $\ell_2$ are used in this product. We focus on two choices for these indices. We define *fully connected* networks, for which $k_1$, $k_2$, $\ell_1$, and $\ell_2$ are allowed to take on all possible values. We also define *simply connected* networks, for which we impose the condition that $k_1 = k_2$ and $\ell_1 = \ell_2$.

The exact choice made in combining these indices affects the dimensionality of the network. The width of the Clebsch-Gordan nonlinearity output in a fully connected network with a maximum spherical degree $L_{\max}$ and a width $d_h$ (i.e., $k \in \{1, ..., d_h\}$) is given by

$$\Omega^{\text{full}}_{\ell, L_{\max}} = d_h^2 \left( \left[ \frac{1}{4}(2 + \ell(5+\ell) - 2\left\lfloor \frac{1}{2}(\ell - 1) \right\rfloor \right] + (\ell+1)(L_{\max} - \ell) \right) + (\ell+1)(L_{\max} - \ell) \tag{S12}$$

Meanwhile, for a simply connected network with the same $L_{\max}$ and $d_h$, the width of the Clebsch-Gordan nonlinearity is

$$\Omega^{\text{simple}}_{\ell, L_{\max}} = d_h (L_{\max} + 1 - \ell) \tag{S13}$$

**Spherical batch normalization.** Batch normalization is a common feature in neural networks that allows for smooth training of the network. Since the output of our nonlinear Clebsch-Gordan product is not bounded, batch normalization becomes extremely important to ensure that activations remain finite throughout training. We impose a batch normalization layer after each linear operation, producing normalized inputs to the nonlinear operation (Clebsch-Gordan product). We use the following batch normalization during training,

$$\hat{F}^{k}_{\ell m} = \frac{\hat{F}^{k}_{\ell m}}{\sqrt{\langle |\hat{F}^{k}_{\ell}|^2 \rangle_{\text{batch}} + \epsilon}} \tag{S14}$$

where $|\hat{F}^{k}_{\ell}|^2 = \sum_{m=-\ell}^{\ell} \hat{F}^{k}_{\ell m} \hat{F}^{*k}_{\ell m}/(2\ell+1)$, $^*$ denotes complex conjugation, $\langle\cdot\rangle_{\text{batch}}$ denotes averaging over a batch, and $\epsilon$ is a small number used to ensure numerical stability, which we set to $\epsilon = 10^{-3}$. During training the network stores a moving average of the norm for each $\ell$ in each channel as

$$N^{\mu,i}_{\ell} = \xi \langle |\hat{F}^{k}_{\ell}|^2 \rangle_{\text{batch}} + (1 - \xi) N^{\mu,i-1}_{\ell} \tag{S15}$$

where $\xi$ serves as the momentum of our moving average, set to $\xi = 0.99$. This moving average is then used as the norm during testing in the following way:

$$\hat{F}^{\mu}_{\ell m} = \frac{\hat{F}^{\mu}_{\ell m}}{\sqrt{N^{\mu,i}_{\ell} + \epsilon}} \tag{S16}$$

The different batch normalization used for training and testing prevents the batching of the validation data to affect the evaluation of the model accuracy during testing.

### III. NETWORK AND TRAINING PROCEDURE FOR H-CNN

**Network architecture.** A complete H-CNN features an equivariant unit of Clebsch-Gordan layer followed by an invariant unit of dense layers as detailed in Fig. S2. A Clebsch-Gordan layer is comprised of an equivariant linearity, a spherical batch norm, a Clebsch-Gordan product, and a concatenation. Invariants from the inputs as well as all outputs of the each CG layer are collected for use in the invariant dense layers (Figs. 1, S2).

**Training procedure.** Networks were trained with all training data being read at run time using the keras `fit` function and a data generator made from a subclass of the `keras.utils.sequeunce` class [16]. Generally 40 workers were used for reading the data and a `max_queue_size` of 900 was found to be optimal for read/evaluation balance. Approximately one tenth of the validation data (51,200 neighborhoods) was evaluated after an approximate one tenth of the training data (256,000 neighborhoods) was seen. This was implemented by using `tf.keras.Model.fit` arguments `steps_per_epoch=1000` and `validation_steps=100`. Data was shuffled at the end of each evaluation interval. These evaluation intervals also served as checkpoints for both early stopping and weight saving. The best network was trained for 8.47 epochs in 4.54 hours on one NVIDIA A40 GPU hosted at the Hyak supercomputer at the University of Washington.

We used stochastic gradient descent and Adam optimizer [17] for training our networks. Adam generally showed better performance, and thus, all networks were trained with Adam during hyperparameter optimization. The main component of the loss was defined as the categorical cross entropy between the one-hot amino acid encoding and the softmax output of the network

$$\mathcal{L} = -\sum_{i=1}^{N} \log p_{\alpha_i} \tag{S17}$$

where $p_{\alpha_i}$ is the network-assigned probability of the true central amino acid $\alpha_i$ in neighborhood $i$.

Some hyperparameters were scheduled according to the behavior of the network's loss. The learning rate was put on a schedule of `ReduceLROnPLateau` to decrease by a factor of 0.1 whenever the validation loss did not improve over 10 epochs, with a minimum learning rate of $10^{-9}$. Ultimately, the networks were trained with early stopping when the validation loss did not improve by a difference of 0.01 over the course of 20 epochs.

**Hyperparameter optimization.** Hyperparameter optimization was performed in two steps: First, we scanned a larger parameter regime and trained 500 networks to identify a reasonable subregion of the parameter space (Fig. S3A). This sampling revealed networks with parameter ranges listed in Table S1. These parameters were then sampled more finely to get the best network models. The resulted training curves are shown in Fig. S3B,C.

It should be noted that we optimized over two classes of networks: (i) fully connected networks, and (ii) the simply-connected networks, as defined in Section II C. In simply connected networks, the nonlinear Clebsch-Gordan products are restricted to the square of Fourier coefficients with the same $\ell$ and channel index, in contrast to the fully connected networks that use bilinear products between all sets of Fourier coefficients (see equation (S11)). These restricted nonlinearities allowed us to use linear layers that had roughly one order of magnitude larger output dimensions in simply vs. fully connected networks (150 as opposed to 20); see Fig. S3E. We separately optimized the hyperparameters for these two network classes.

**Model performance.** For both the simply connected and the fully connected networks, we find the best hyperparameters that minimize the validation loss function in equation S17. The best simply-connected model had 62% classification accuracy and a minimal validation loss of 1.2 while the best fully-connected model had 68% classification accuracy and a minimal validation loss of 0.91. This difference in predictive power of the models appeared consistent across all training scenarios.

We compare the accuracy and the training efficiency of our model to the existing methods that classify amino acids based on all-atom representations of their local environments in Table S2. For this comparison, we used the metric of testing accuracy of the amino acid class since this was more commonly reported in the literature. Specifically, we compare H-CNN with two 3D CNN methods that used conventional convolutional neural networks on voxelized protein structures [18, 19]. Notably, these 3D CNN methods require local alignment of neighborhoods to the central amino acid. Also, each of these methods uses a cube of side length 20Å, while spheres of radius 10Å are used as inputs to H-CNN. Our method thus sees approximately half ($\pi/6$) of the volume that these other models see. Overall, our model has a comparable accuracy to the best 3D CNN method [19], but it is much more efficient in training.

We also compare our model to other methods that prioritize rotational symmetry, i.e., the Spherical CNN introduced in ref. [20] and the 3D Steerable CNN from ref. [21]. Spherical CNN [20] is approximately rotationally invariant by performing 3D convolutions on a discretized grid over the 3D sphere. 3D Steerable CNN [21] is a rotationally invariant method by applying spherical convolutions in a steerable basis. H-CNN performs better than both of these methods in the classification task with an order of magnitude fewer parameters (Table S2).

**Alternative models.** We studied the effect of our post-processing of atomic neighborhoods by performing ablation studies on charge and SASA. For each ablation, we reran the second stage of our hyperparameter optimization and only considered fully connected networks. The overall accuracy from removing SASA was 57% while the overall accuracy from removing charge was 56%, in contrast to the 68% accuracy of the complete model shown in Fig. 2. A similar 10% drop in accuracy associated with SASA and charge was previously reported in ref. [19]. Fig. S4 shows the confusion matrices for each model compared to the best model's confusion matrix.

### IV. COMPARING H-CNN WITH EVOLUTIONARY MODELS FOR AMINO ACID PREFERENCES

As shown in the main text, the interchangeability of amino acids predicted by H-CNN is consistent with the evolutionary substitution patterns reflected by the BLOSUM62 matrix (Fig. 2D). In addition to this coarse-grained measure, we also assessed the consistency of substitutions at the single-site level between the H-CNN predictions and evolutionary data. Evolutionary variation at a given site in a protein reflects both the substitutability of amino acids at that position (e.g. due to their common physico-chemical properties) and the epistatic interactions with the other covarying residues in the protein.

To isolate the site-specific effects of substitutions, we used the software EV-Couplings [22] to infer a model for amino acid preferences (and their couplings), using amino acid covariations in multiple sequence alignments (MSAs) of protein families. Specifically, EV-Couplings infers a maximum likelihood Potts model for interactions between residues in a protein sequence [23]. From this model, we can evaluate the relative preference for occurrence of different amino acids at a given position in a protein sequence.

We used EV-Couplings to infer models of amino acid preferences for all of the protein domains in our test set. For each protein chain in our test data, we used EV-Couplings to compute the MSA for the protein associated with UniProt ID listed under the associated PDB entry. The MSA is computed using UniRef90 [24] that clusters homologous proteins such that sequences within each cluster have at least 90% identity to and 80% overlap with the longest sequence (seed). For inference of the Potts model, we used the default EV-Couplings settings for the rest of the parameters. Ultimately we were able to infer evolutionary models on 67 protein families corresponding to 67 chains in our testing set. These protein families amounted to 11,221 unique sites, for which we compare the EV-Couplings predictions on amino acid preferences with the H-CNN predictions in Fig. 2F.

To measure the similarity between the predictions of H-CNN and EV Couplings, we define the profile overlap $\rho$ for amino acid preferences at a given position. Specifically, for two sets of amino acid probabilities $P_A, P_B \in [\mathbb{R}^+_{\leq 1}]^{19}$, the vector overlap is defined as the normalized dot-product between the centered probability vectors,

$$\rho = \frac{\big(P_A - \langle P_A \rangle_{\text{sites}}\big) \cdot \big(P_B - \langle P_B \rangle_{\text{sites}}\big)}{\sqrt{|P_A - \langle P_A \rangle_{\text{sites}}|^2 |P_B - \langle P_B \rangle_{\text{sites}}|^2}} \tag{S18}$$

where $\langle \cdot \rangle_{\text{sites}}$ denotes the average of probability vector overs all sites in the data. Vector overlap takes values between -1 and 1, with 1 indicating a perfect alignment between the two (centered) vectors of amino acid preferences, and -1 indicating complete disagreement.

### V. EFFECT OF SHEAR PERTURBATION ON PROTEIN MICRO-ENVIRONMENTS

We locally perturbed protein structures around their crystal conformation with the shearing deformation. As demonstrated in Fig. 3, shearing by an angle $\delta$ at a given residue of a protein is a perturbation to the backbone dihedral angles at two neighboring residues given by

$$(\phi_i, \psi_{i-1}) \longrightarrow (\phi_i + \delta, \psi_{i-1} - \delta) \tag{S19}$$

This perturbation was chosen because it can substantially change the local protein structure near the residue of interest while minimally affecting the far-away residues.

Physical interactions in a protein should be dependent on the pairwise distances between atoms. We define $D_{ab}$ to be the pairwise distance in angstroms between a pair of atoms $(a, b)$ in the protein structure. Then $\Delta D^{i,\delta}_{ab} = D^{i,\delta}_{ab} - D^0_{ab}$ measures the deviation of these pairwise distances from the pairwise distances in the native (unperturbed) structure $D^0_{ab}$, when site $i$ is sheared by an angle $\delta$ (Fig. 3). We estimate the total shear-induced distortion in a protein structure as the root-mean-square of the deviations of pairwise distances $\text{RMS}\Delta D^{k,\delta}_{ij}$; here, the average is taken over all pairs of atoms that were at distances smaller than 10 Å in the unperturbed structure, i.e., $\{(a, b) \,|\, D^0_{ab} < 10\,\text{Å}\}$.

We characterize the model's response to the perturbation by the change in the protein network energy (Fig. 3), defined as the sum of pseudo-energies (logits) of amino acids over all sites in the protein,

$$E = \sum_{i \in \text{sites}} \tilde{E}^{\alpha_i}_i \tag{S20}$$

where $\alpha_i$ denotes the protein's amino acid at site $i$.

As shown in Fig. 3D, the native protein structure appears to be at the energy minimum of the network and shear perturbations often result in less favorable protein micro-environments. To assess the robustness of this result, we evaluated the protein network energy in eq. S20 for alternative amino acids, while keeping the surrounding protein structure intact (i.e., using the wild-type neighborhood for model evaluation). Fig. S5 shows that if amino acid substitutions preserve the physico-chemical properties of the residue (i.e., substitutions according to the BLOSUM62 matrix), the undistorted structure remains close to the energy minimum, yet the landscape becomes shallower. However, for random amino acid substitutions, no energy minima is recovered. These results further suggest that H-CNN has learned an effective physical potential for protein micro-environments.

[9] Schrödinger, LLC. The PyMOL Molecular Graphics System, Version 1.8; 2015.

[10] Gallagher T, et al. Two Crystal Structures of the B1 Immunoglobulin-Binding Domain of Streptococcal Protein G and Comparison with NMR. Biochemistry. 1994;33(15):4721–4729. doi:10.1021/bi00181a032.

[11] Batzner S, et al. E(3)-Equivariant Graph Neural Networks for Data-Efficient and Accurate Interatomic Potentials. Nature Communications. 2022;13(1):2453. doi:10.1038/s41467-022-29939-5.

[12] Tung WK. Group theory in physics. Philadelphia: World Scientific; 1985.

| Hyperparameter | variable | broad sampling space | sampling scheme | fine sampling space | optimal fc network value | optimal sc network value |
|---|---|---|---|---|---|---|
| batch size | $n_b$ | [8, 5000] | uniform | 256 | 256 | 256 |
| dropout rate | $\lambda_{dr}$ | [0,0.6] | uniform | $[0, 10^{-5}]$ | $5.49 \times 10^{-4}$ | $1.95 \times 10^{-2}$ |
| hidden dimension | $d_h$ | fc:[6,20], sc:[50,160] | uniform | fc:[8,20], sc:[100,500] | 14 | 109 |
| No. CG layers | $n_l$ | [1,5] | uniform | [4,5] | 4 | 5 |
| No. dense layers | $n_d$ | [1,5] | uniform | [2,4] | 2 | 2 |
| Max spherical degree | $L_{max}$ | [2,7] | uniform | [4,5] | 5 | 5 |
| learning rate | $\lambda_l$ | $[10^{-6}, 10^{-1}]$ | exponential | $10^{-3}$ | $10^{-3}$ | $10^{-3}$ |
| regularization strength | $\lambda_r$ | $[10^{-6}, 10^{-1}]$ | exponential | $[10^{-10}, 10^{-16}]$ | $1.2 \times 10^{-16}$ | $1.89 \times 10^{-14}$ |

TABLE S1. **Hyperparameter bounds and optimal values.** Hyperparameters varied during model optimization are listed. The two different bounds used during the two steps of hyperparameter optimization are shown in the broad and fine sampling spaces. Different bounds for fully connected (fc) and simply connected (sc) networks are shown when appropriate. Finally, the optimal value is listed for both the optimal fully connected and simply connected networks.

| method | rotationally invariant | dataset | post-processing | N | No. of parameters | training time | accuracy |
|---|---|---|---|---|---|---|---|
| H-CNN | yes | ProteinNet | charge<br>hydrogen<br>SASA | $2.8 \times 10^6$ | $3.6 \times 10^6$ | 4.54 hours | 68% |
| 3D CNN [18] | no | SCOP & ASTRAL | None | $7.2 \times 10^5$ | $10^7$ | 3 days | 40% |
| 3D CNN [19] | no | SCOP & ASTRAL | PDB-REDO[25]<br>charge<br>hydrogen<br>SASA | $1.6 \times 10^6$ | $6.1 \times 10^7$ | – | 70% |
| Spherical CNN [20] | approx. | PISCES | charge<br>hydrogen | – | $6 \times 10^7$ | – | 56% |
| Steerable CNN [21] | yes | SCOP & ASTRAL | PDB-REDO<br>charge<br>hydrogen<br>SASA | $1.6 \times 10^6$ | $3.3 \times 10^7$ | – | 58% |

TABLE S2. **Comparison of structure-informed models for protein neighborhoods.** H-CNN and existing methods trained on classifying all-atom neighborhoods are listed along with the summary quantities of the models. H-CNN demonstrates the power of respecting symmetry since it has fewer parameters and trains faster than 3D-CNNs despite being trained on at least the same amount of data.

| variant | I3Y | I3P | V87M | D92N | R96H | R96N | R96D | R96W | R96Y | L99G | M102K | M102L | M106K | M120K | V149M | V149S | V149C | V149G | G156D | T157I |
|---|---|---|---|---|---|---|---|---|---|---|---|---|---|---|---|---|---|---|---|---|
| PDB id | 1L18 | 1L97 | 1CU3 | 1L55 | 1L34 | 3CDT | 3C8Q | 3FI5 | 3C80 | 1QUD | 1L54 | 1L77 | 231L | 232L | 1CV6 | 1G06 | 1G07 | 1G0P | 1L16 | 1L10 |
| $\Delta\Delta G$ [kcal/mol] | -2.3 | | -2.3 | -1.4 | | -3.0 | -3.5 | -4.5 | -4.7 | -6.3 | -6.9 | -2.11 | -3.4 | -1.6 | -2.8 | -4.4 | -2.0 | -4.9 | -2.3 | -1.2 |
| pH | 6.5 | | 5.4 | 5.7 -5.9 | | 5.35 | 5.35 | 5.35 | 5.35 | 5.4 | 5.3 | 5.7 | 3 | 3 | 5.4 | 5.4 | 5.4 | 5.4 | 6.5 | 6 |
| source | [26] | [27] | [28] | [29] | [30] | [31] | [31] | [31] | [31] | [32] | [33] | [34] | [35] | [35] | [28] | [36] | [36] | [36] | [37] | [38] |

| variant | I3V | T26S | S38N | S38D | G77A | A82P | A93T | M106I | M106L | E108V | T109N | T109D | N116D | S117F | M120Y | M120L | N144D | V149I | T151S | T152V |
|---|---|---|---|---|---|---|---|---|---|---|---|---|---|---|---|---|---|---|---|---|
| PDB id | 1L17 | 131L | 1L61 | 1L19 | 1L23 | 1L24 | 129L | 1P46 | 234L | 1QUG | 1L59 | 1L62 | 1L57 | 1TLA | 1P6Y | 233L | 1L20 | 1G0Q | 130L | 1G0L |
| $\Delta\Delta G$ [kcal/mol] | -0.4 | | 0.6 | | 0.4 | 0.8 | | 0.2 | 0.5 | 0.7 | 0.1 | 0.6 | 0.6 | 1.1 | -0.1 | 0.5 | | -0.1 | | 0.2 |
| pH | 6.5 | 5.4 | 5.7 -5.9 | 5 | 6.5 | 6.5 | 5.4 | 5 | 3 | 5.4 | 5.7 -5.9 | 5.7 -5.9 | 5.7 -5.9 | 5.4 | 5 | 3 | 5 | 5.4 | 5.4 | 5.4 |
| source | [26] | [39] | [29] | [40] | [41] | [41] | [39] | [42] | [35] | [32] | [29] | [29] | [29] | [43] | [42] | [35] | [40] | [36] | [39] | [36] |

TABLE S3. **T4 lysozyme variants.** Each of the 40 variants, shown in Fig. 4, with the respective protein structure is listed along with the corresponding PDB entry. When available, the $\Delta\Delta G$ associated with a mutation and the pH of the experimental setup in which protein stability was measured are provided.

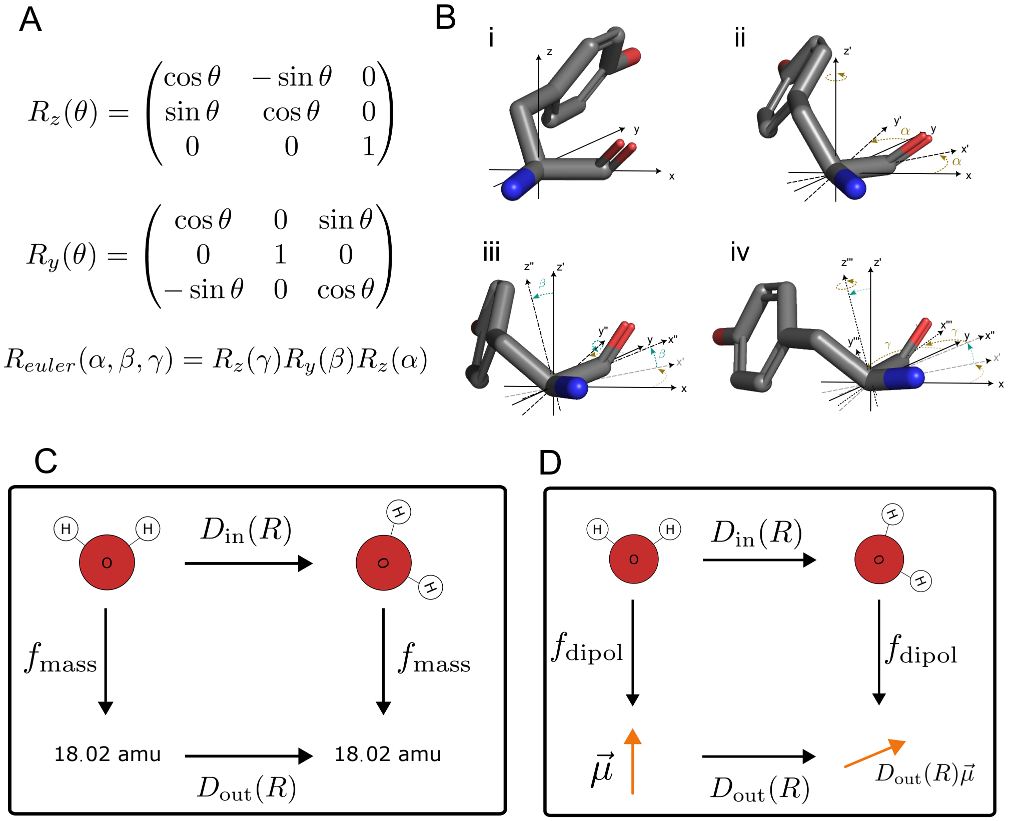


Fig. S1. **Parameterization of rotations and rotational equivariance.** We use x-y-z Euler angles to parameterize rotations in 3D. This parametrization **(A)** can be written in a matrix form, and **(B)** it can be visualized on a 3D object. Starting from a given orientation in (B.i), the object's rotation can be broken down into three steps: First, rotation around the z-axis of angle $\alpha$ (B.ii). Second, rotation around the new y-axis by angle $\beta$ (B.iii). And finally, rotation around the new z-axis by angle $\gamma$ (B.iv). A map from the geometric descriptions of an object to physical observables can be invariant or equivariant under rotations. **(C)** A rotationally invariant map, such as $f_{\text{mass}}$ which gives the mass of a molecule, remains unchanged under the molecule's rotation. **(D)** A rotationally equivariant map, such as $f_{\text{dipol}}$ which gives the dipole moment of a molecule, transforms in a well-defined way with the molecule's rotation, denoted by the operator $D_{\text{in}}(R)$. Specifically, in this case, the transformation operator for the dipole moment under rotation $D_{\text{out}}(R)$ acts linearly on the dipole moment in the original frame and is uniquely determined by the parameters of the rotation $R$.

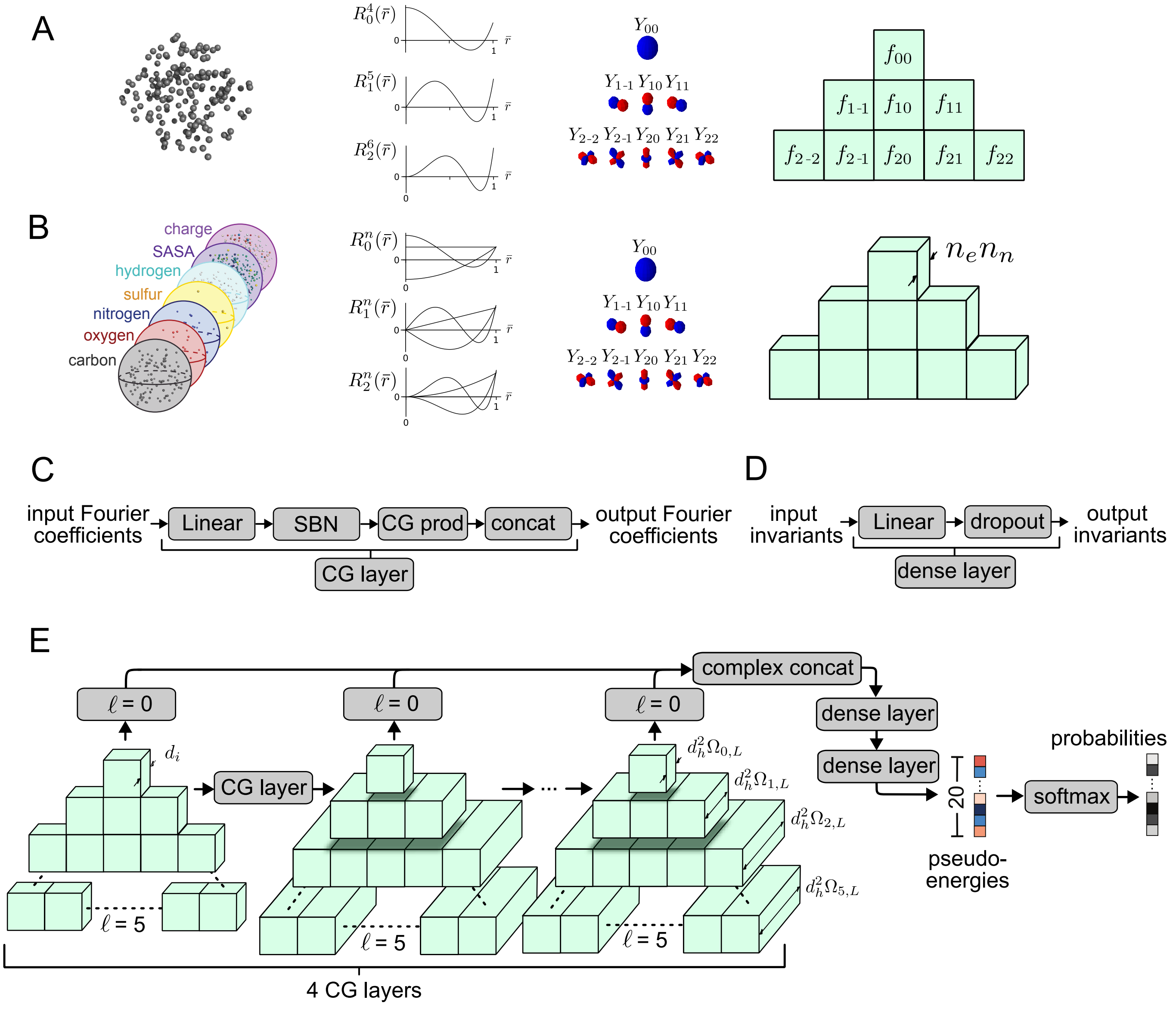


Fig. S2. **Schematics of rotationally equivariant data encoding and network architecture in H-CNN. (A)** Any point cloud (left) can be written in the equivariant spherical harmonic Fourier basis by integrating the point cloud density against the spherical harmonics along with a choice of a radial function (middle), which depends on the spherical harmonic degree $\ell$. The triangular structure (right) demonstrates the resulting Fourier coefficients $f_{\ell m}$, in which the rows reflect the different degrees of spherical harmonics $0 \leq \ell$, which are non-negative integers. Individual cells within each row correspond to different orders $m$ of the spherical harmonics, which are integers bounded by $-\ell \leq m \leq \ell$. **(B)** The Fourier transform becomes three dimensional when radial resolution is required thereby increasing the number of discrete $n$-values used $n_n$. The multiplicity of coefficients is also determined by the number of point clouds being transformed $n_\rho$ (e.g. the different atomic channels on the left). Since both the point clouds and the radial information are invariant under rotations, these two dimensions can be combined in index to give preference in organizing information according to the degree $\ell$ and order $m$ of the spherical harmonics. **(C)** The Clebsch-Gordan block (CG block) is fully equivariant and is comprised of a linear layer, a spherical batch norm (SBN), Clebsch-Gordan product (CG prod), and a degree-wise concatenation (concat). **(D)** Throughout all layers of the network, all invariants ($\ell = 0$) are collected and fed into a series of dense layers which are simple linear combinations with training dropout. **(E)** These two main components in (C,D) comprise the bulk of the entire network. First the input Fourier coefficients are processed through successive CG layers. Invariants are collected from the original input and from the output of every CG layer. The real and imaginary components of these invariants are split and concatenated (complex concat) and then fed into a series of dense layers. This schematic reflects the dimensions of the optimal model discovered in this work. Four CG layers were used. We note that the maximal width of the network as determined by the output of the nonlinearity depends on $\ell$ due to the selection rules of the CG prod. We denote this width $d_h^2\Omega_{\ell,L}$ where $L$ is the maximal degree $\ell$ used in the network and are given in eqs. (S12) and (S13). Two dense layers were used in the optimal model. The dimensions of these layers are $(4852 \times 500)$ and $(500 \times 20)$. The output of these dense layers are termed pseudo-energies and act as logits in a softmax to estimate probabilities for each amino acid. We emphasize the only nonlinearities in the network are the Clebsch-Gordan products and the ultimate softmax.

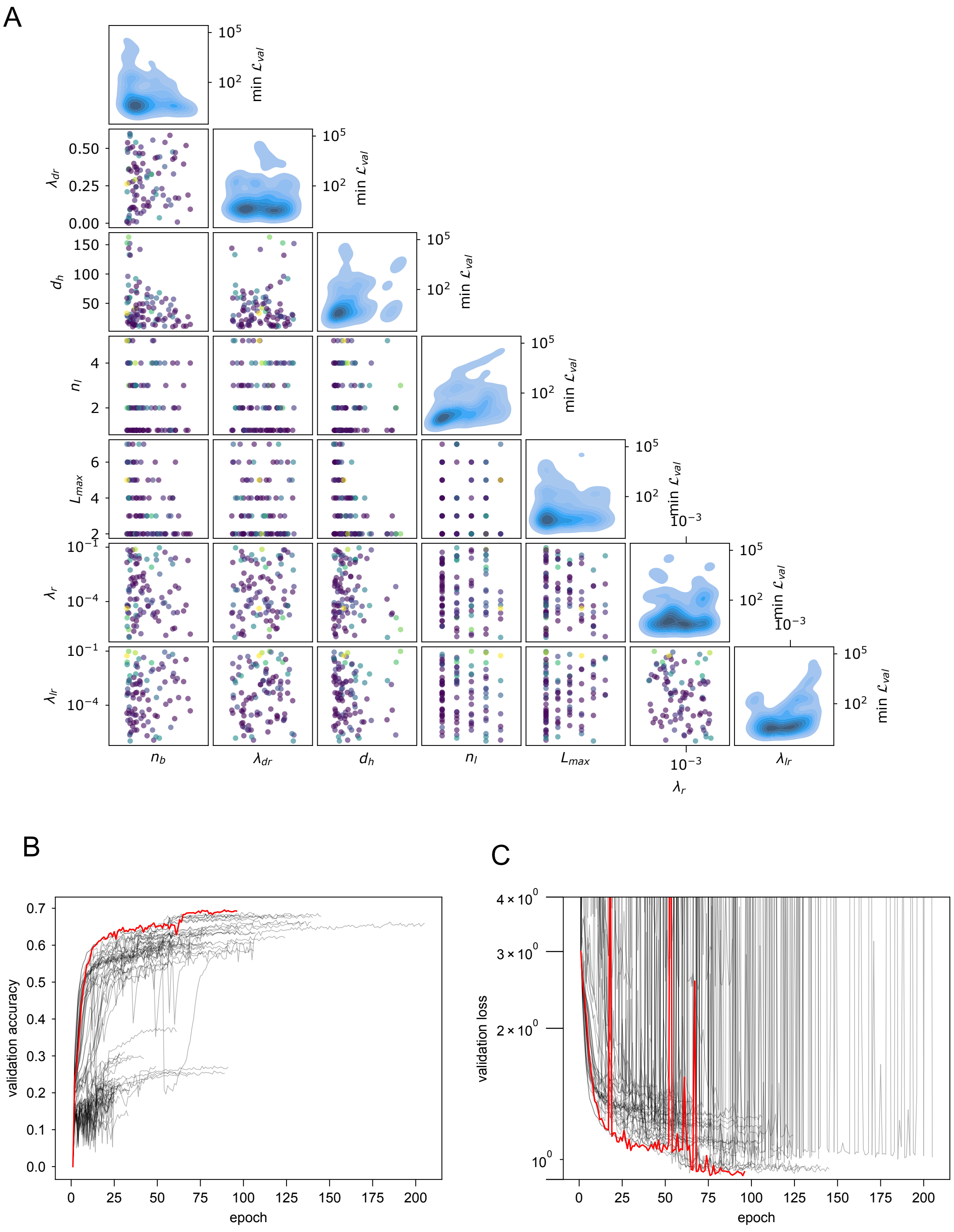


Fig. S3. **Hyperparameter optimization for H-CNN.** Hyperparameter optimization was performed in two steps for both the simply connected and the fully connected networks: First, hyperparameters were scanned across a wide regime of hyper-parameters via random sampling. From this scan, a narrow band of hyper-parameters were analyzed to determine the optimal network **(A)**. Off the diagonal, each dot's color shows the validation loss seen during this first step of hyperparameter tuning for a network with the combination of hyperparameters shown on the two axes. On the diagonal, the density of networks is shown as a function of loss and the hyperparameter tested. Validation accuracy **(B)** and loss **(C)** are shown as a function of training epochs for all networks trained in the second stage of hyperparameter tuning. Red lines show the network with the best validation loss that is used throughout this work.

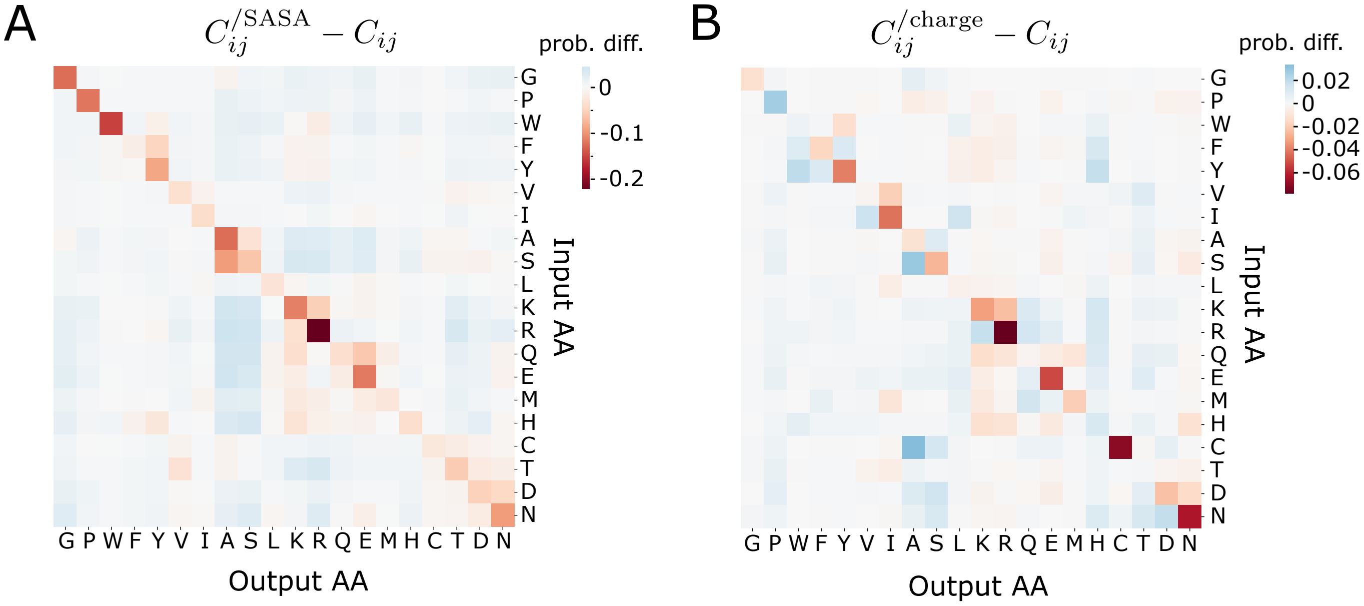


Fig. S4. **H-CNN performance after charge and SASA ablation.** The difference in the confusion matrix for networks with ablated **(A)** SASA and **(B)** charge with respect to the network with the full information (Fig. 2A) is shown. Negative (red) values demonstrate less probability assigned to the input/predicted pair when the input is ablated, while positive (blue) values show increased probability mass under ablation. The removal of SASA (A) appears to decrease the network's ability to predict all amino acids as seen on the diagonal. The effect is most pronounced for hydrophobic amino acids in the upper left, with some hydrophilic amino acids (R, K, E) also displaying strong effects. Interestingly, most of the probability mass is on average redistributed to small amino acids A and S. When charge is removed (B), the network demonstrates worse predictions on charged and polar amino acids most notably R, C, N, and E. The overall network's accuracy after removing SASA and Charge are 57% and 56%, respectively.

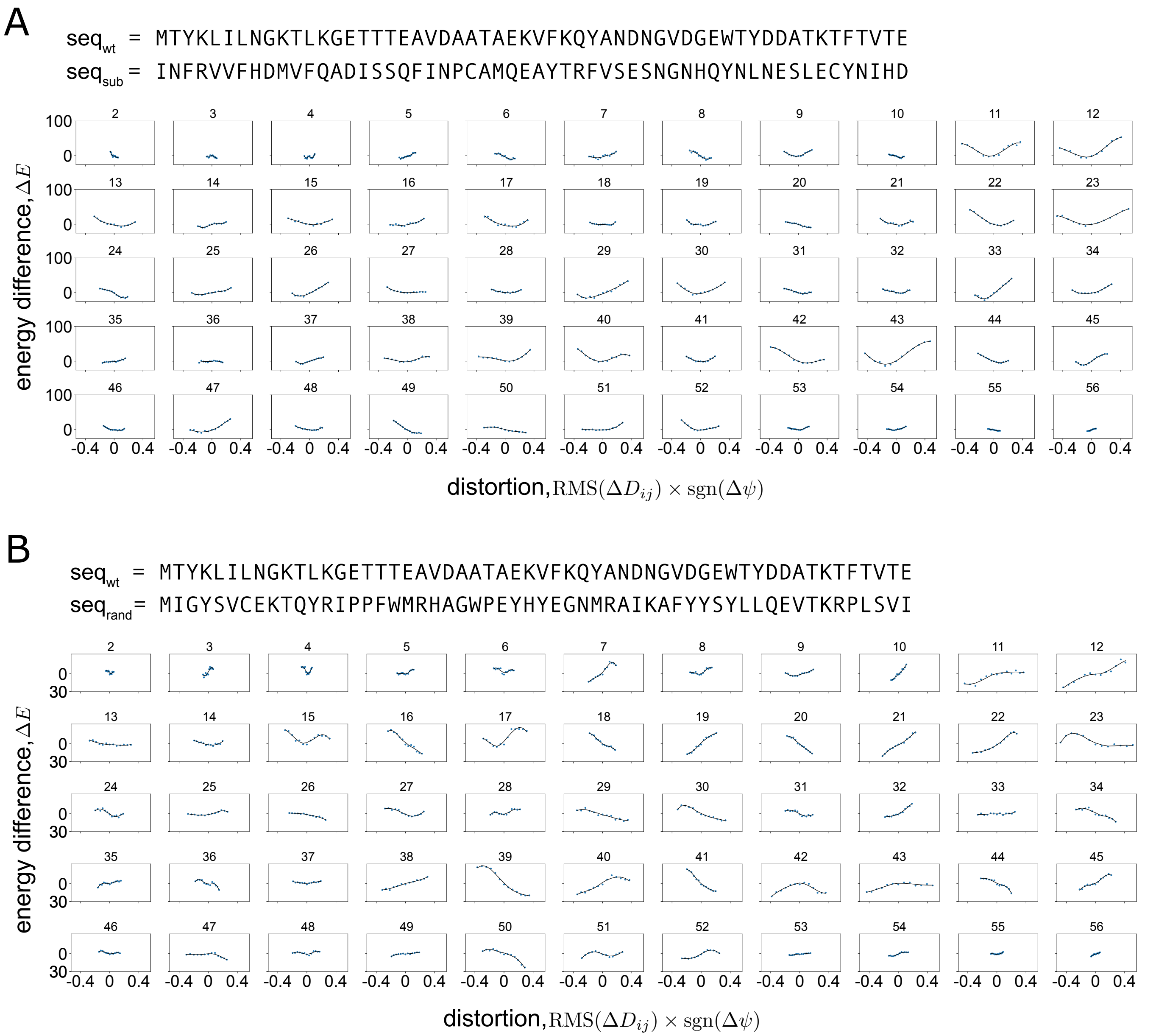


Fig. S5. **Robustness of response to shear perturbation.** Similar to Fig. 3, but the change in network energy is evaluated for alternative amino acids at the shear position, while keeping the surrounding protein structure intact (i.e., using the wild-type neighborhood for model evaluation). The alternative amino acids are drawn according to the BLOSUM62 matrix in **(A)**, and randomly in **(B)**; the wild-type and the alternative sequence are shown each panel. When amino acids are substituted according to their BLOSUM62 scores, potential wells are generally recovered (A), while no energy minima is recovered for random substitutions (B). Specific sequences used are given by $\mathrm{seq}_{\mathrm{sub}}$ (substitutions according to the BLOSUM62 matrix), and $\mathrm{seq}_{\mathrm{rand}}$ (random substitutions) in each panel and are shown in comparison to the wildtype sequence $\mathrm{seq}_{\mathrm{wt}}$.

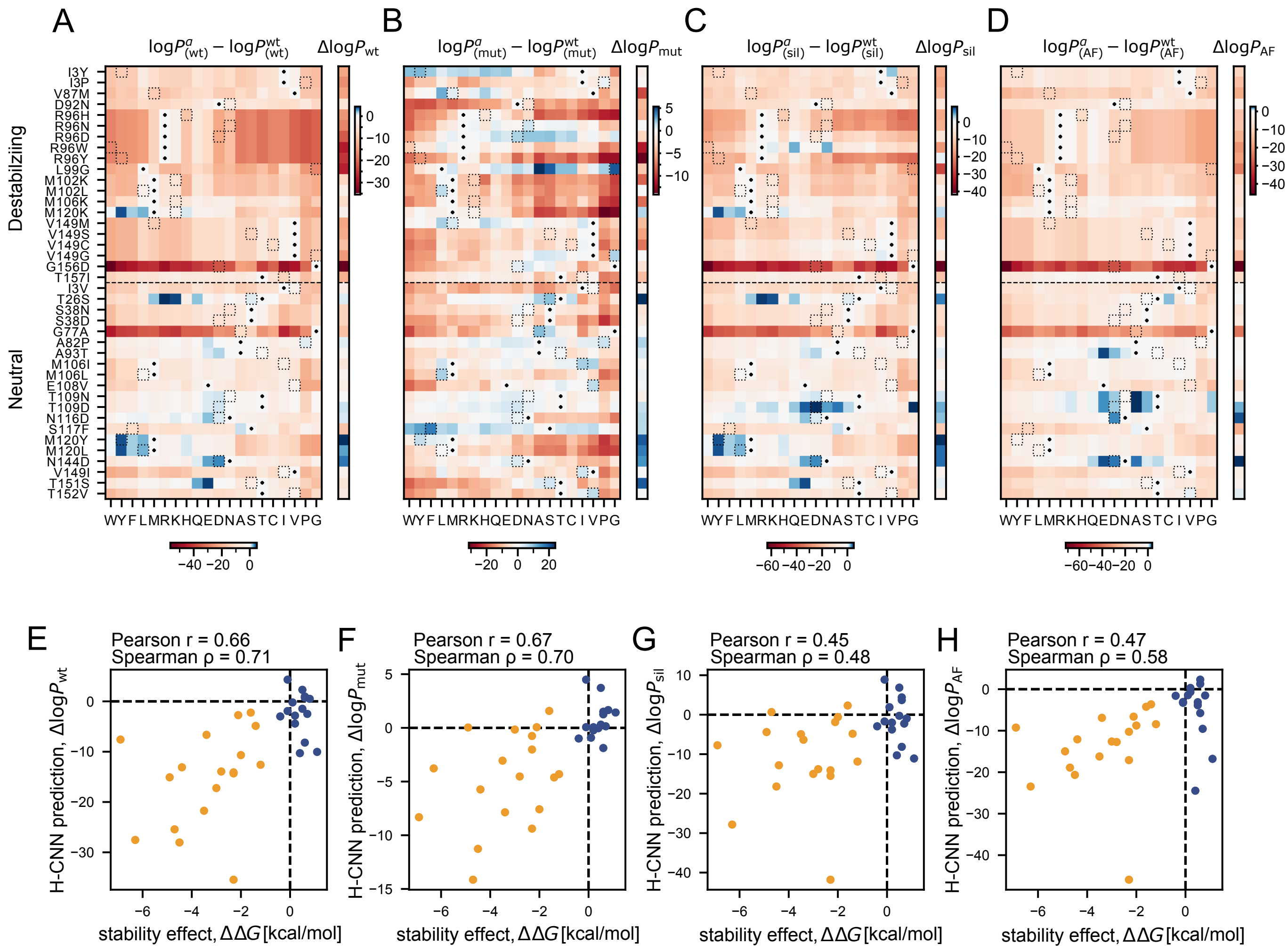


Fig. S6. **Predictions for the stability effect of mutations in T4 lysozyme with different protein structures. (A-D)** The large heatmaps show the H-CNN predicted log-probability of different amino acids relative to the wild-type at the substituted positions along the T4 lysozyme, indicated on the left-hand side of panel A (similar to Fig. 4A). In each heatmap, the dots indicate the identity of the wildtype amino acid, and the dotted squares indicate the amino acid in the specified position for the mutated variant in each row. The predictions for log-probabilities are evaluated using the wild-type structure (A), the mutant crystal structures (B), the wild-type structure relaxed *in-silico* via PyRosetta for the specified substitution (C), and the computationally predicted structure of the wild-type T4 lysozyme using AlphaFold (D). The relative log-probabilities of the mutations of interest are shown separately in one column next to each heatmap. They are calculated with the structural information that is available in different scenarios. For the wild-type, mutant, and *in silico* structures, $\Delta \log P_* = \log P^{\text{mut}}_{(*)}/P^{\text{wt}}_{(\text{wt})}$, with $*$ indicating the specified protein structure in each case. This definition is akin to $\Delta\Delta G$. For the AlphaFold structure, we use $\Delta \log P_{\text{AF}} = \log P^{\text{mut}}_{(\text{AF})}/P^{\text{wt}}_{(\text{AF})}$—a quantity which could be evaluated in the absence of any experimental structural knowledge for a protein. **(E-H)** The predicted relative log-probabilities shown in panels (A-D) are plotted against the experimentally measured $\Delta\Delta G$ values for each variant. Destabilizing mutations are shown in orange and neutral/beneficial mutations are shown in blue. Overall, H-CNN predictions correlate well with the mutational effects on protein stability, with the Pearson and Spearman correlations indicated in each panel and AUC values to discriminate between destabilizing and neutral mutations indicated in Fig. 4D. Accounting for changes in the local protein structures due to mutations in (B, C) sets the correct reference point in predictions such that the estimated log-probability ratio is overall positive for neutral/beneficial mutations and negative for destabilizing mutations.

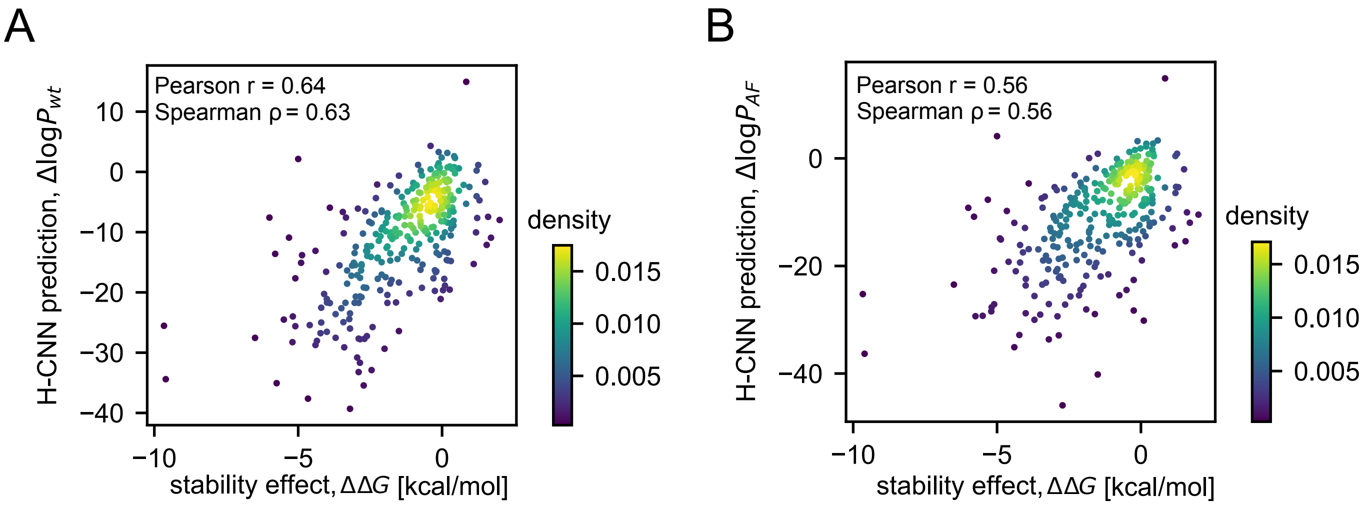


Fig. S7. **H-CNN predictions for the stability effect of all available single point mutations in T4 lysozyme.** H-CNN predictions for the relative log-probabilities using **(A)** the wild-type structure $\Delta \log P_{\mathrm{wt}}$ and **(B)** the AlphaFold predicted structure $\Delta \log P_{\mathrm{AF}}$ are shown against the experimentally measured $\Delta\Delta G$ values for 310 single point mutation variants of T4 lysozyme. Mean $\Delta\Delta G$ was used when multiple experiments reported values for the same variant. The colors show the density of points in each panel as calculated via Gaussian kernel density estimation. H-CNN predictions correlate well with the experimental values, with the Pearson and Spearman correlations indicated in each panel.

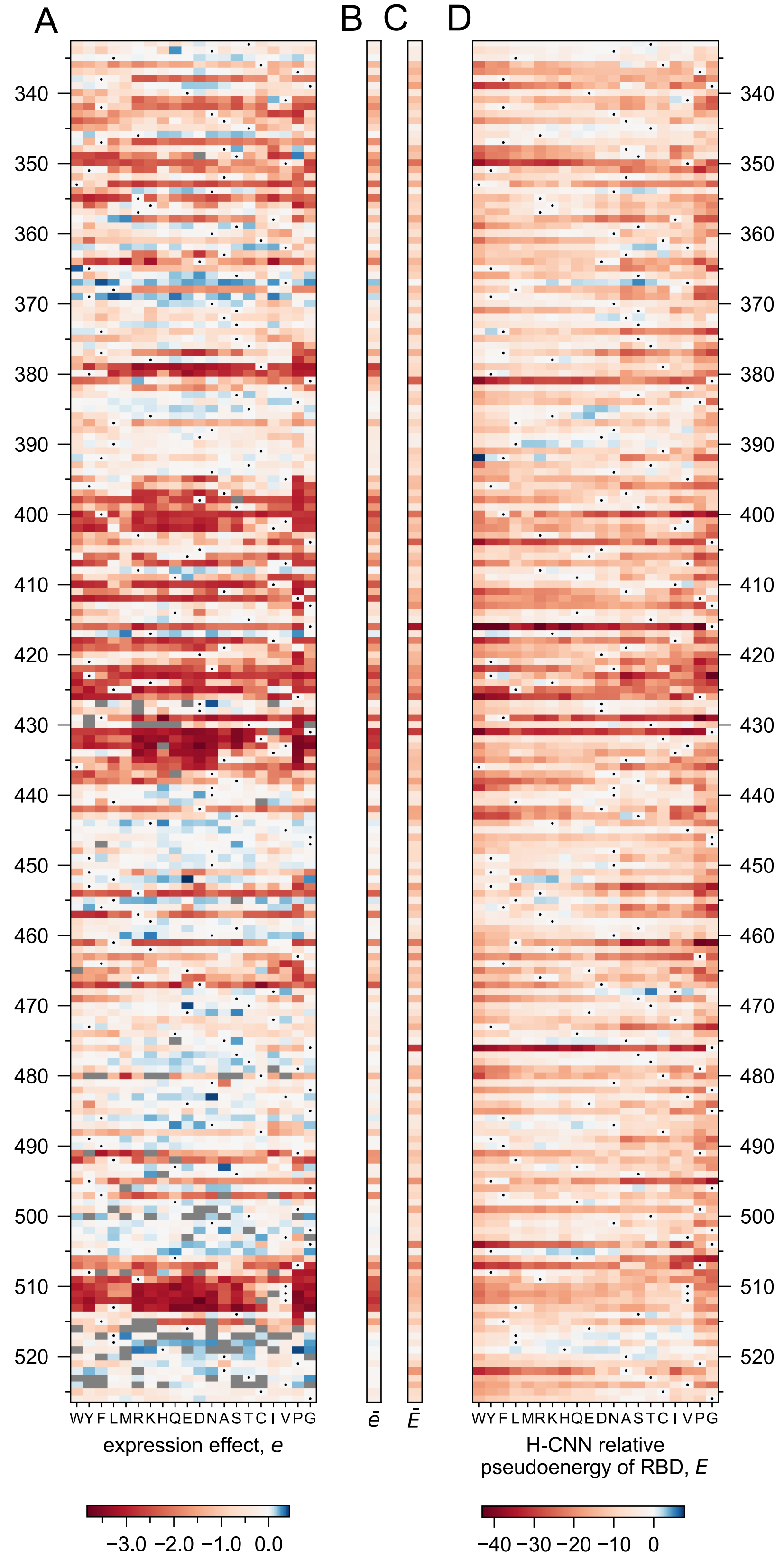


Fig. S8. **Experimental RBD expression of SARS-CoV-2 in DMS experiments and the H-CNN predictions. (A)** Experimental expression of single-point mutation variants of SARS-CoV2 RBD relative to the wildtype (dots) as measured using yeast surface display experiments. Expression effect $e$ is calculated as $\Delta\log_{10}(\text{MFI})$ where MFI is the mean fluorescent intensity. Mutations to G or P dominate the negative tail of expression effects. To better show the effect of all mutations, the negative portion of the colorbar is bounded by the expression of the most detrimental variant that is not a mutation to G or P. **(B)** The mean expression effect of mutations $\bar{e}$ is listed next to each site. **(C,D)** The H-CNN predicted stability effect of mutations (D), and the mean predicted stability effect for each site $\bar{E}$ (C), evaluated based on the computationally isolated RBD structure, are shown. Again the negative portion of the colorbar is bounded from below by the most destabilizing prediction not involving mutations to G or P.

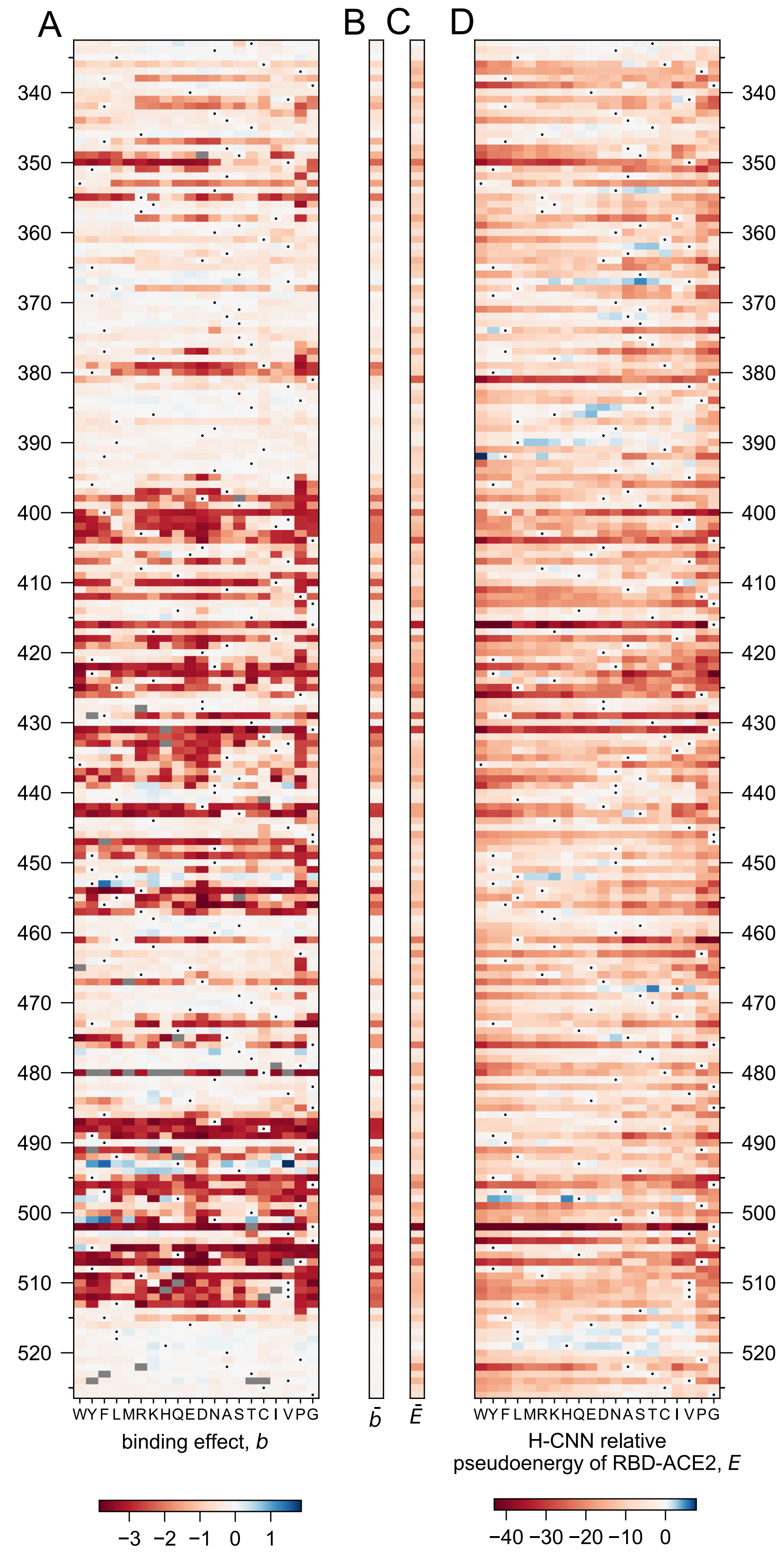


Fig. S9. **Experimental RBD-ACE2 binding affinity in DMS experiments and the H-CNN predictions. (A)** Experimental binding of single-point mutation variants of SARS-CoV2 RBD to the human SCE2 receptor relative to the wild-type (dots) as measured using yeast surface display experiments. Binding effect $b$ is reported from $\Delta \log K_D$ where negative values represent weaker binding relative to the wildtype. Mutations to G or P dominate the negative tail of expression effects. To better show the effect of all mutations, the negative portion of the colorbar is bounded by the expression of the most detrimental variant that is not a mutation to G or P. **(B)** The mean binding effect of mutations $\bar{b}$ is listed next to each site. **(C,D)** The H-CNN predicted effect of mutations (D) and the mean predicted stability effect for each site $\bar{E}$ (C), evaluated based on the crystal structure of RBD-ACE2 complex, are shown. Again the negative portion of the colorbar is bounded from below by the most destabilizing prediction not involving mutations to G or P.

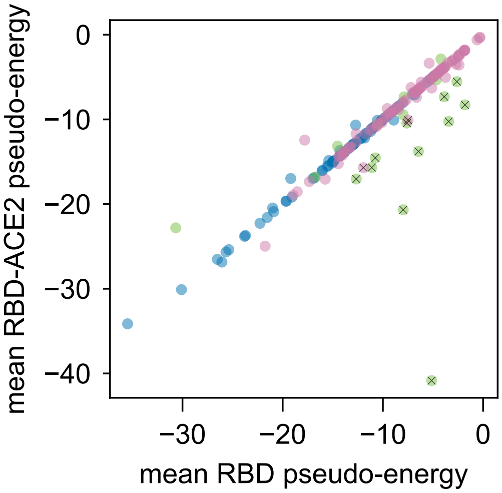


Fig. S10. **H-CNN predictions for stability and binding of RBD.** The H-CNN predicted mean pseudo-energy per site predicted from the RBD-ACE2 protein complex (measure of binding) is plotted as a function of the mean pseudo-energy per site, inferred from the isolated RBD structure (measure of stability). Points are colored according to their experimental values of expression and binding as illustrated in Fig. 5. Most points lie on the diagonal since the RBD pseudo-energy is determined from the RBD-ACE2 crystal structure with the ACE2 masked. However, sites at the RBD-ACE2 interface show deviations from the diagonal. H-CNN prediction for sites that are tolerant of mutations for stability but not binding are indicated by crosses (below the diagonal points with large stability values). These sites tend to be at the RBD-ACE2 interface and overlap with the green category from Fig. 5.
